# The human blood harbors a phageome which differs in Crohn’s disease

**DOI:** 10.1101/2024.06.04.597176

**Authors:** Quentin Lamy-Besnier, Ilias Theodorou, Maud Billaud, Hao Zhang, Loïc Brot, Antoine Culot, Guillaume Abriat, Harry Sokol, Marianne De Paepe, Marie-Agnès Petit, Luisa De Sordi

## Abstract

Increasing evidence suggests that the human blood hosts microbes including bacteria and eukaryotic viruses, which could have important implications for health. Bacteriophages of the blood are challenging to study and have been overlooked, but could translocate to this environment from different body-sites. We thus developed specific virome protocols and analysis methods to study the viral communities of blood samples obtained from healthy individuals and Crohn’s disease (CD) patients. We uncovered a diverse viral community in the human blood, dominated by phages infecting Pseudomonadota bacteria. We found that an important fraction of those phages overlaps with the gut virome, consolidating the idea that gut phages can translocate to the blood. Strikingly, viral communities of the blood were different between CD patients and healthy individuals, revealing a new signature of disease. This was not the case for fecal viral communities. Collectively, these results advance our knowledge of the microorganisms present in the human blood and pave the way for further studies of this environment in the context of disease.

## Introduction

Despite being initially thought to be sterile, increasing evidence shows that the human blood contains bacteria^1–5^, although they do not form a common community across individuals^6^. The blood also hosts viruses, but efforts have been mainly directed at identifying eukaryotic viruses, overlooking bacterial viruses, bacteriophages (phages)^7–10^. Phages are important members of human-associated microbiota, particularly in the gastrointestinal tract (GIT) where they have been linked to a variety of diseases^11–14^. Their entry into the blood has also been suggested by studies showing the translocation of phages across intestinal cell layers *in vitro*^15,16^. Therefore, it is possible that a community of phages exists in the blood and could be relevant to human health. Recently, the sequencing of circulating DNA in a human cohort of sepsis revealed a blood phageome that could be employed to identify the pathogens causing the infection^17^. These findings underscore the significant potential of blood phage communities, and call for further studies characterizing the encapsidated and circulating viral particles of the human blood phageome to fully leverage its possibilities as a marker of disease.

The limited number of studies on blood microbes can be explained by very low concentrations of microorganisms, which is a challenge for typical metagenomic experimental and analytical approaches. The issue is exacerbated for viruses having small genomes, like phages, resulting in small quantities of total DNA that are hardly compatible with metagenomic sequencing approaches. Contamination of the samples by environmental DNA is also a particular concern in this context. A keystone study used over 8,000 human blood samples to identify multiple viruses, including prevalent human herpesviruses and phages, but could not discriminate contamination from reliable signals^18^. This study and most others used only total DNA, as opposed to performing a size filtration to remove bacteria from the samples (virome sequencing), which led to reduced relative abundances of viruses. Novel and virus-focused studies are thus required for the reliable and accurate exploration of the human blood virome.

Crohn’s disease (CD) is a chronic, inflammatory bowel disease characterized by recurrent inflammation of the GIT. Despite ever-increasing worldwide incidence, no specific marker of this complex disease has been identified^19^. On top of genetic and environmental causes, the gut microbiota composition has been strongly associated with CD^20–22^. However, this signature becomes blurred when the analysis is restricted to the viral fraction of the microbiota. Initially, the gut virome was found to be correlated with CD^23–25^, but nowadays it is no longer considered linked with the disease due to its intrinsically high interindividual variability^26^. This variability is driven by the broad diversity of the gut virome. Therefore, in order to investigate the potential link between the human virome and CD, other viral communities with a reduced complexity might be more appropriate.

Here, we developed a specific experimental and bioinformatic approach to address the challenges of the blood environment in order to characterize the human blood virome. Using this method on a cohort comprised of healthy and CD individuals, we first show that the blood virome is essentially composed of a diverse community of phages infecting mostly bacteria of the Pseudomonadota phylum (previously known as Proteobacteria). In order to explore the origins of these phages, we took advantage of fecal samples obtained from the same individuals, as well as of databases of gut and oral viruses. We found that the gut virome was an important source of blood phages. Remarkably, the comparison of healthy and CD individuals uncovered a difference in the blood virome composition, whereas no such distinction was observed in the fecal virome. Among the differentially abundant phages, all of those that are less abundant in CD patients infect the genus *Acinetobacter*, which marks a first step towards unravelling the connection between CD and the blood virome. These data provide the first comprehensive overview of the human blood phageome and underline its potential to better understand complex diseases such as CD.

## Results

### Accurate determination of vOTUs in blood and fecal samples

To study their viral communities, we collected blood and fecal samples from 15 CD patients and 14 healthy individuals (**Fig. 1A** and **Table S1**). Since blood samples contain low amounts of DNA and are prone to contamination^18,27^, we set up two negative controls consisting of a blood collection tube and a standard microtube, both filled with sterile double-distilled water before processing. Viral DNA was enriched from all samples, prepared and sequenced (Methods and **Table S2**). We developed a specific pipeline for the analysis of these samples (**Fig. 1B-C**). After cleaning and removing the human reads, an assembly was performed for each individual, as well as using all the samples together (cross-assembly), leading to a total of 165,933 contigs larger than 2 kb. The contigs were clustered at the species level, reducing the number of operational taxonomic units (OTUs) to 85,985. To select viral OTUs (vOTUs), we used a combination of dedicated tools and viral databases (Methods), detecting a total of 17,454 vOTUs (**Fig. S1A-B-C**). The quality of vOTUs was assessed using CheckV^28^ (**Fig. S2A-B**). These vOTUs were then characterized through annotation, host prediction and classification. The sequence and annotation of all vOTUs are publicly available, and complete information for each vOTU can be found in **Table S3**.

**Figure 1:**
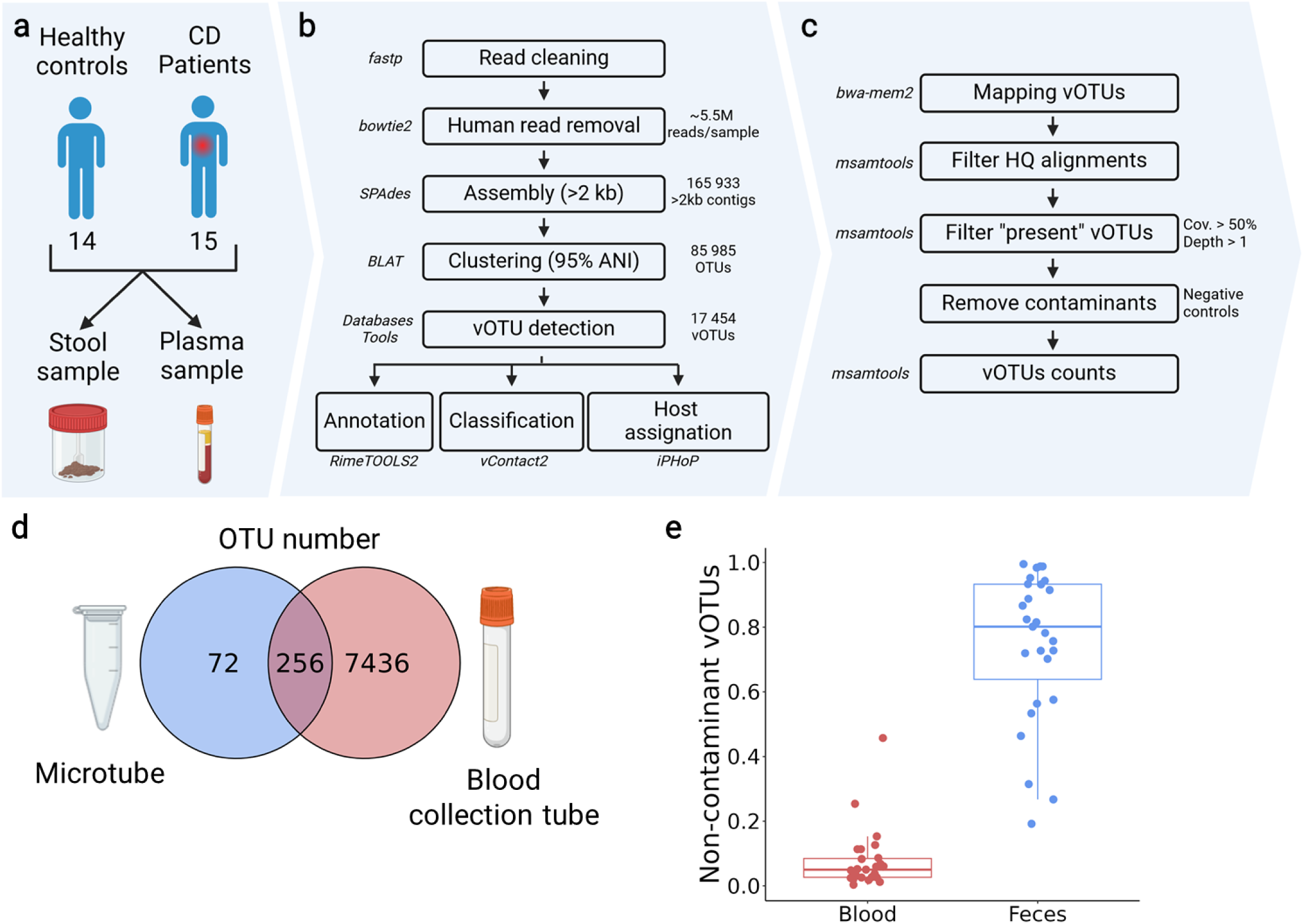
Determination of the human blood virome. **a.** Schematics of the cohort and sample collection. **b.** Bioinformatic pipeline for the detection of vOTUs. **c.** Bioinformatic pipeline for the determination of vOTU counts. **d.** Number of OTUs in the two negative control samples. **e.** Relative abundance of the non-contaminant (i.e. not found in the negative control samples) vOTUs per sample type and disease status (orange = CD patients, green = healthy individuals).

The blood being an environment characterized by a low abundance of viruses, we used a stringent filter to only consider OTUs for which sufficient evidence for their presence in each sample was achieved. These criteria were such that on average, each nucleotide of the OTU must be sequenced at least once (depth ≥ 1), and at least 50% of the OTU should be sequenced (coverage ≥ 50%). Only the OTUs passing these criteria were considered as present in a given sample. On average, this led to the removal of 80% OTUs per sample, which had few reads mapped to them but not sufficiently (**Fig. S3A**).

Next, to identify and remove contaminant OTUs from our samples, the content of negative control samples was determined. This analysis revealed a high number of contaminants (total of 7,764 OTUs), and a surprising difference between the microtube and the blood collection tube, with the latter being much richer in contaminants (**Fig. 1D**). This highlights the importance of the adequate choice of controls in metagenomics studies. When looking at the classification of these OTUs, bacterial contamination dominated (6,047 OTUs), followed by viral (484 OTUs) and eukaryotic (438 OTUs) sequences (**Fig. S4A**). Both tubes were heavily contaminated with DNA from microbes associated with the skin microbiota: bacteria belonging to the Actinomycetota phylum, including families such as *Proprionibacteria*, as well as phages predicted to infect such hosts, and yeasts from the genus *Malassezia* (**Fig. S4B-C-D)**. This suggests that microorganisms from the human skin are responsible for an important part of the contamination. A core of contaminant sequences was found across the blood samples, showing that the contaminants are not tube-specific and were appropriately identified using our two negative controls (**Fig. S5**). We favoured a stringent approach and removed all the OTUs present in any of these two negative controls. This filtering step led to the loss of an average of 57% of the reads for the blood samples, and 4% for the fecal samples (**Fig. S3B**).

Viromes usually contain a portion of non-viral DNA, which in our case represented on average 83% of the reads in blood samples and 23% in fecal samples (**Fig. S3C**). This fraction consisted mostly of bacterial DNA (**Fig. S6A-B-C**) and was also removed. Finally, non-contaminant vOTUs represented around 6% of the reads for the blood (**Fig. 1E**), highlighting the importance of the filtering steps. As expected, these filters had a small impact on the fecal samples, with an average of 75% of reads per sample kept.

By designing an adequate and conservative pipeline, we thus have access to the viral composition of each sample with high confidence.

### The human blood virome is diverse and dominated by phages infecting Pseudomonadota

Blood samples carried a median of 132 vOTUs, for a total of 1,794 unique vOTUs present across the 27 samples, demonstrating that the human blood virome contains a diverse viral community (**Fig. 2A**). The spiking of human samples with two phages, SPP1 (dsDNA) and M13 (ssDNA), led to estimating that around 10^5^ viral particles are present per mL of blood (**Fig. 2B**). This is the first attempt at quantifying the number of viral particles in the blood virome. Altogether, these results converge on the presence of a relatively large community of viruses in the blood.

**Figure 2:**
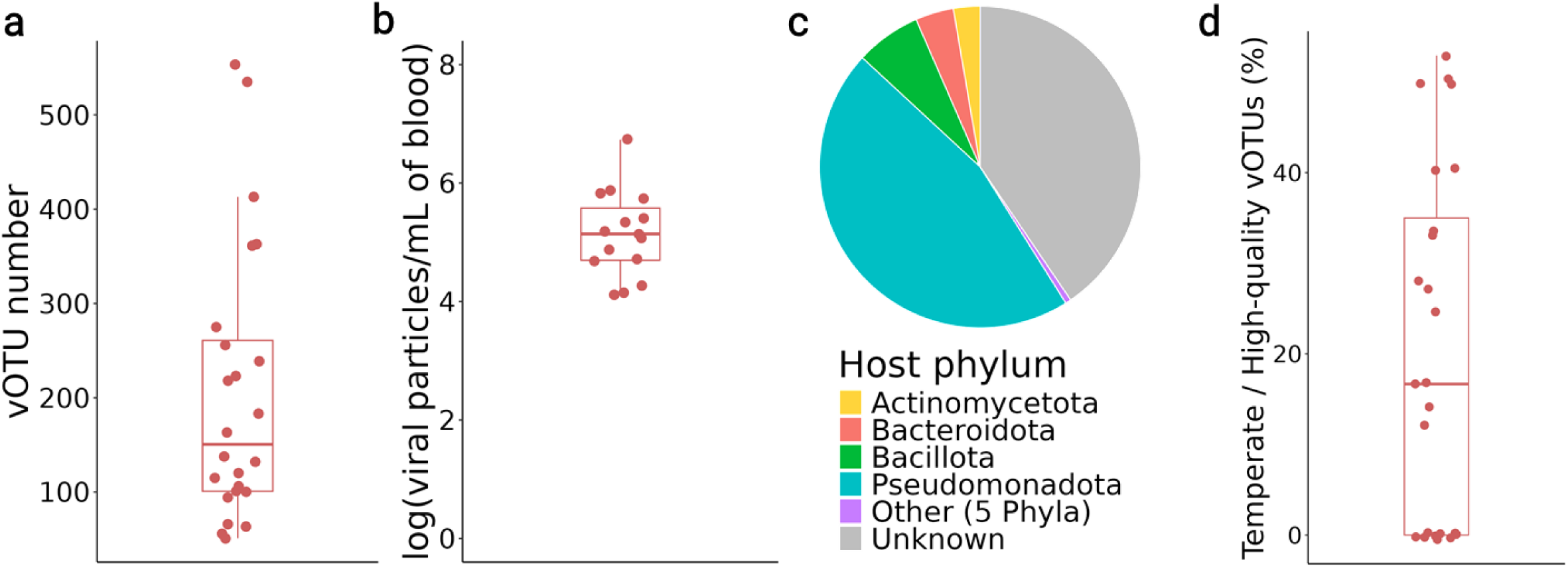
Overview of the human blood virome. **a.** Number of vOTUs per blood sample. **b.** Estimated number of viral particles per mL of blood and per blood sample. **c.** Distribution of the predicted host phyla for all the vOTUs found in blood samples. **d.** Percentage of vOTUs with a temperate lifestyle among the high-quality vOTUs. The temperate lifestyle was predicted by the presence of genes coding for an integrase, and only the high-quality vOTUs were considered (Methods).

We then characterized the nature of this human blood virome. Firstly, we were able to detect eukaryotic viruses previously reported to be frequent in human blood, mainly Anelloviruses^7,10^ (**Fig. S7**). Interestingly, Herpesviruses, which are also frequently reported in the human blood^8,9,18^ were found in some blood samples but also in the negative controls, so their natural presence in this ecosystem cannot be ascertained. However, phages largely dominated the viral community of the human blood. This phageome was mostly composed of phages infecting Pseudomonadota bacteria (> 80% relative abundance), followed by phages infecting bacteria from the phyla Bacillota (formerly known as Firmicutes), Bacteroidota (formerly Bacteroidetes) and Actinomycetota (**Fig. 2C**). This is in line with the main bacterial phyla found in the human blood^1–3,5^, as well as with other human-associated microbiota. Another significant characteristic of viral communities is their lifestyles, which can be classified as either virulent, where viruses lyse their host to release progeny, or temperate, where viruses can integrate into the host chromosome as prophages and replicate passively. By looking at the presence of integrase genes in high-quality or complete vOTUs, we estimated that around ∼20% of the blood high-quality vOTUs were temperate (**Fig. 2D**), and they collectively represented 20% of the high quality vOTUs abundances (**Fig. S8A**). Taking these well-assembled vOTUs as a proxy for the whole community, we predict that temperate phages should represent collectively ∼20% of the blood viral community.

Collectively, we uncover a diverse viral community in the human blood, dominated by phages infecting mostly Pseudomonadota bacteria.

### Origins of the human blood virome

Two main hypotheses could explain the presence of phages in the human blood: induction of prophages integrated in blood bacteria, or translocation from other body-sites. As most phages were classified as virulent (**Fig. 2D**), induction does not seem prevalent, which prompted us to explore the second option. Bacteria are able to translocate through the intestinal epithelium in both physiological and disease conditions^29^, and reports have also shown that phages translocate through human cells *in vitro*^16^. The intestinal microbiota also harbors the highest virus density, making the gut virome the most probable origin of blood phages.

It is to investigate this hypothesis that we sequenced fecal samples collected from the same individuals, and at the same time as the blood samples. Their analysis revealed typical characteristics of the human gut virome, which illustrates the reliability of our analysis pipeline. In particular, we found a median of 722 vOTUs per fecal sample (**Fig. S9A**), for a total of 10^8^-10^9^ viral particles per gram of feces (**Fig. S9B**), with predicted infected hosts being mainly Bacteroidota and Bacillota bacteria (**Fig. S9C**), and an estimation of around 50% of temperate phages based on high-quality vOTUs (**Fig. S9D** and **Fig. S8B**). When comparing the vOTUs present in the blood to those present in the fecal samples, only ∼3% of blood vOTUs overlapped, representing ∼1.5% of vOTUs per sample and a corresponding ∼1.5% of abundance (**Fig. 3A-B** and **Fig. S10A**). A detailed study of these shared vOTUs is included in a dedicated, parallel article where we looked at the nature of viruses that could translocate from the gut to the bloodstream ^30^. To complement this analysis, we broadened the search for gut phages in the blood virome by comparing our blood vOTUs with recently released databases of gut viruses ^31–35^. This resulted in ∼20% of blood vOTUs having homology (>90% identity over >90% target length) with gut phages (**Fig. 3C**), representing ∼20% of the relative abundance of blood viruses (**Fig. S10B**). This suggests that the gut is indeed an important source for the blood virome, although it does not account for the majority of blood phages. It also illustrates the value of leveraging large databases built from samples collected at multiple sections of the GIT, as opposed to our limited number of samples of fecal origin.

**Figure 3:**
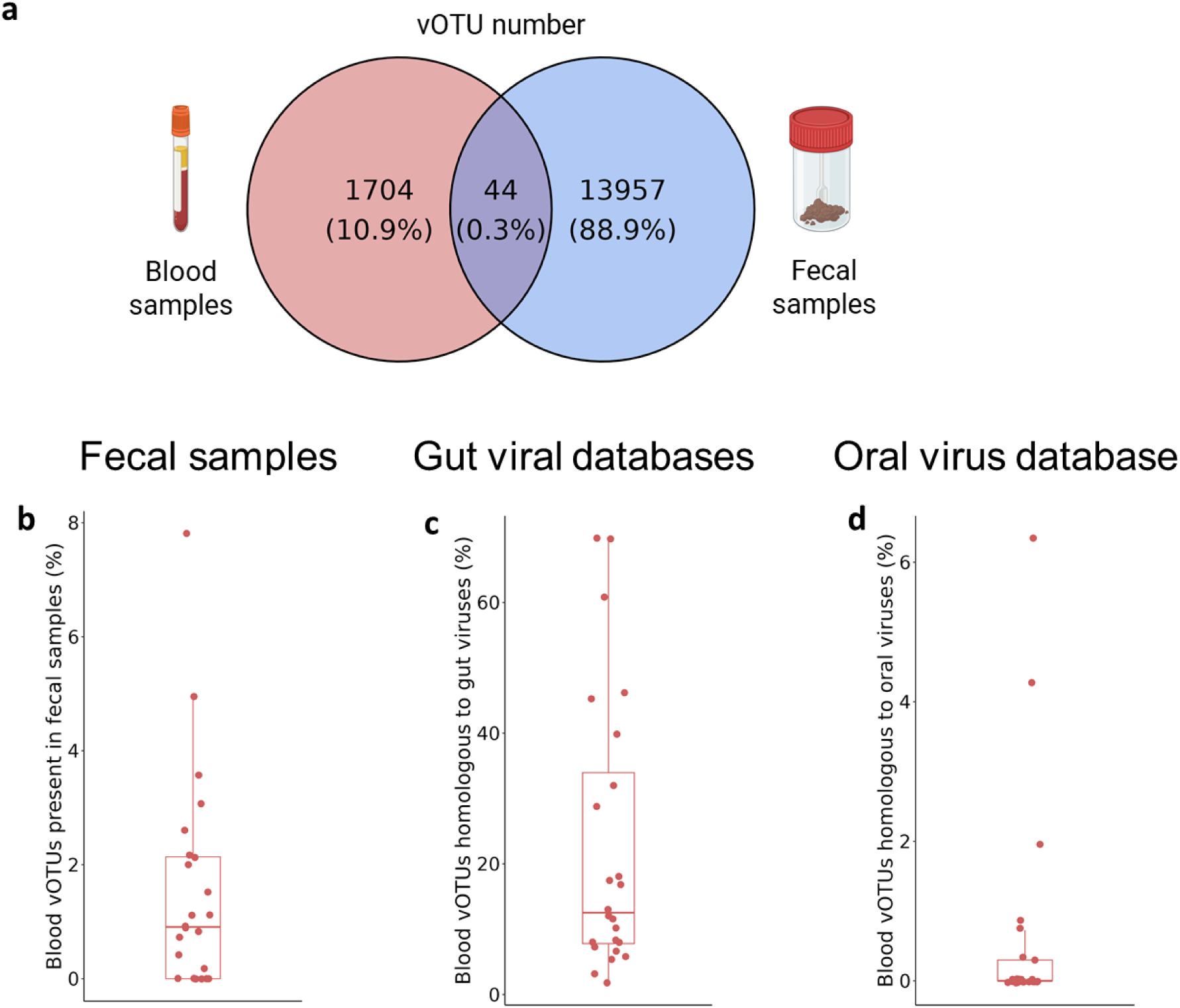
The origins of the human blood virome. **a.** Venn diagram of the vOTUs identified in blood and fecal samples. **b.** Percentage per blood sample of vOTUs that are also found in fecal samples. **c.** Percentage per blood sample of vOTUs that have a close homolog in a gut viral database. **d.** Percentage per blood sample of vOTUs that are have a close homolog in the Oral Virus Database.

In an attempt to find the origin of the remaining 80% blood viruses, we interrogated the database of viruses found at another body-site from which phages could translocate: the mouth. The oral microbiota contains an important fraction of Pseudomonadota bacteria^36^. As this phylum hosts most of our blood phages, the mouth appears to be relevant candidate. We thus compared the blood vOTUs with viruses from the Oral Virus Database^37^. We found ∼2% of blood vOTUs with high similarity to oral phages (>90% identity over >90% target length), which together represented ∼2% of relative abundance (**Fig. 3D** and **Fig. S10C**), showing that the oral virome also contributes to the blood virome.

Altogether, while the origins of the blood virome are difficult to establish, the present analysis suggests that the gut virome is an important contributor.

### The blood virome differs in Crohn’s disease

We then proceeded to compare the blood viral community between healthy individuals and CD patients (**Fig. 4A**). Most of the metrics presented previously, such as the proportion of temperate phages or the number of viral particles per mL, did not differ between the two groups (**Fig. S11A-B**). Instead, we found a difference in the host phyla that the vOTU are predicted infect: CD blood viromes presented an increase in phages infecting Bacteroidota (p=0.03) (**Fig. S11C**). This phylum is similarly increased in the gut microbiota of CD patients^38^, suggesting that changes in the blood virome may be linked to those in the gut bacteriome.

**Figure 4:**
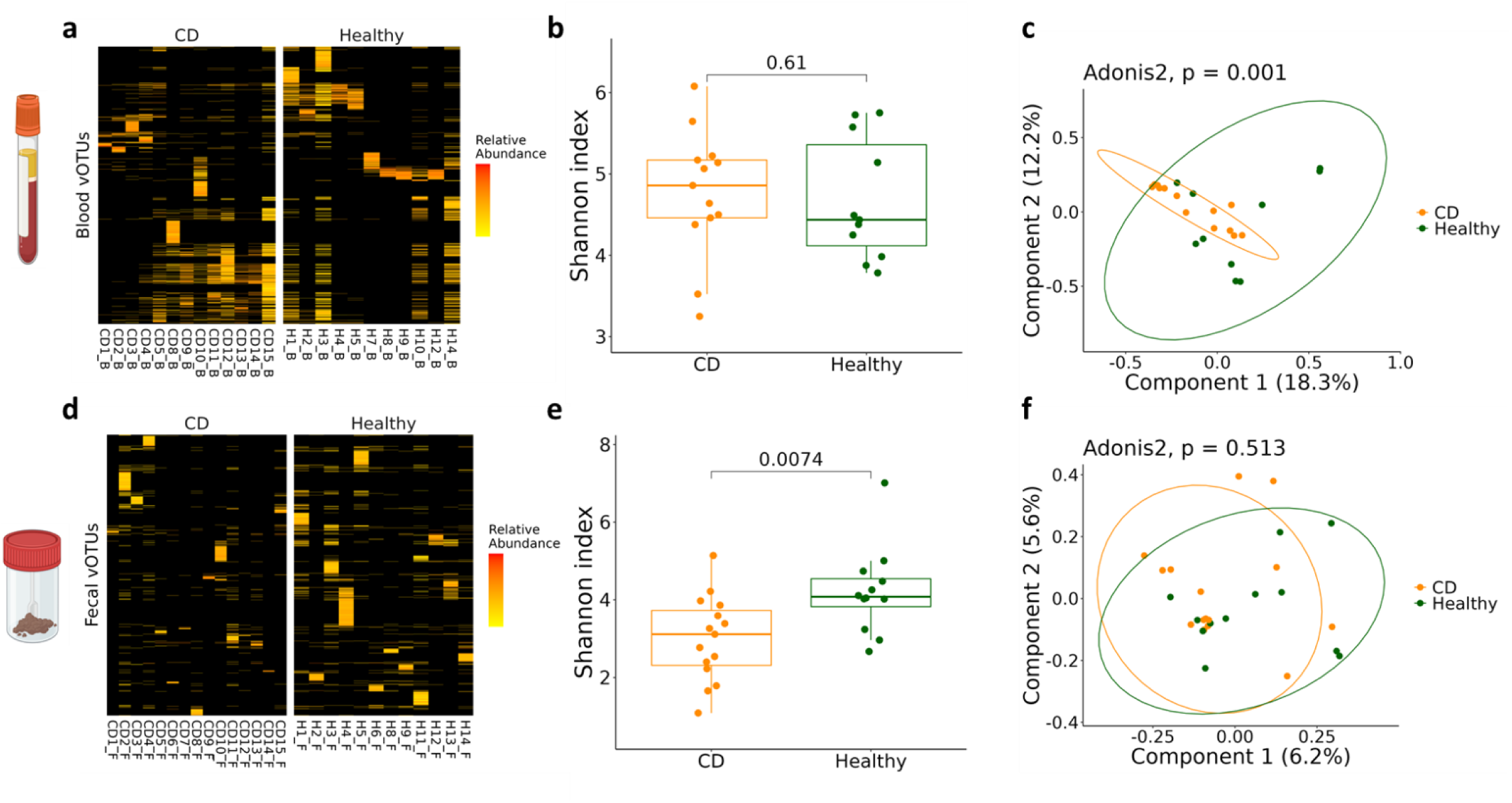
Differences between healthy individuals and CD patients in blood and fecal communities. **a.** Heatmap of the relative abundance of blood vOTUs across blood samples, separated by disease status. **b.** Alpha diversity of the blood samples measured by the Shannon index and separated by disease status. **c.** PCoA of the vOTU composition of the blood samples, using the Bray-Curtis dissimilarity distances. The associated Adonis2 analysis revealed a significant difference between both groups (p=0.001). **d.** Heatmap of the relative abundance of fecal vOTUs across fecal samples, separated by disease status. **e.** Alpha diversity of the fecal samples measured by the Shannon index and separated by disease status. **f.** PCoA of the vOTU composition of the fecal samples, using the Bray-Curtis dissimilarity distances. The associated Adonis2 analysis revealed no significant difference between both groups (p=0.513).

We also compared the alpha-diversity between both groups, measured by the Shannon index, and did not find any significant differences (**Fig. 4B**). Similar observations were made with other diversity metrics, including richness as well as Simpson and Chao1 indexes (**Fig. S12A-B-C**). However, comparing the composition of both communities through an analysis of beta-diversity revealed a strong difference between the two groups (Bray-Curtis dissimilarity, p=0.001, **Fig. 4C**). While the healthy group displayed essentially a scattered distribution, the CD blood samples clustered together. These results were confirmed using the Jaccard index (p=0.001, **Fig. S12D**) and indicate that the blood virome harbors a signature of CD.

Furthermore, when the same analysis was performed with fecal samples (**Fig. 4D**), we did not observe any major changes in the viral communities (**Fig. S11D-E-F**). One of the main shifts observed in the gut microbiota of CD patients is a reduction of the species *Faecalibacterium prausnitzii*^20,21^. We saw a comparable diminution of the relative abundance of phages infecting the *Faecalibacterium* genus in CD patients (p=0.012, **Fig. S11G**), suggesting similar changes in the gut bacteriome and virome. More generally, a slight decrease in viral diversity in the fecal samples of CD patients was identified (p=0.0074), which could be due to the reduced bacterial diversity associated with CD (**Fig. 4E** and **Fig. S12E-F-G**). Importantly, the two groups are not separated in terms of beta-diversity (**Fig. 4F** and **Fig. S12H**). This can be explained by inter-individual variability of the gut virome which has been shown to hinder attempts at finding disease signals with metagenomic approaches^26^.

Taken together, our results suggest that the blood virome is more discriminant for CD than the gut virome.

### *Acinetobacter*-infecting phages are depleted in CD individuals

In order to uncover which phages could be responsible for this different blood viral community in CD, we then performed a differential analysis. A total of 46 differentially abundant vOTUs were identified (**Fig. 5**), which consolidates the difference between the blood viral communities of healthy and diseased individuals. In contrast, the same analysis run on the fecal samples did not reveal any differentially abundant vOTUs, being in line with the lack of difference observed by beta-diversity metrics. Among the differentially abundant blood vOTUs, 27 were enriched in CD patients, while 19 showed increased abundance in healthy individuals. While the 27 vOTUs more abundant in CD patients had a variety of predicted hosts, with mostly Pseudomonadota and few Bacillota bacteria, a striking pattern emerged for the 19 vOTUs more abundant in healthy individuals, which were all infecting bacteria from the genus *Acinetobacter*. These results suggest that the changes of the blood virome associated with CD have a precise effect on the healthy blood virome, reducing the population of phages infecting *Acinetobacter* while promoting the presence of a variety of other phages without a clear pattern. Around 15-20% of the differentially abundant phages were temperate, for both the over- and under-represented vOTUs (using vOTUs of all qualities in this case), showing that prophage induction does not explain the changes in the blood virome of the CD patients.

**Figure 5:**
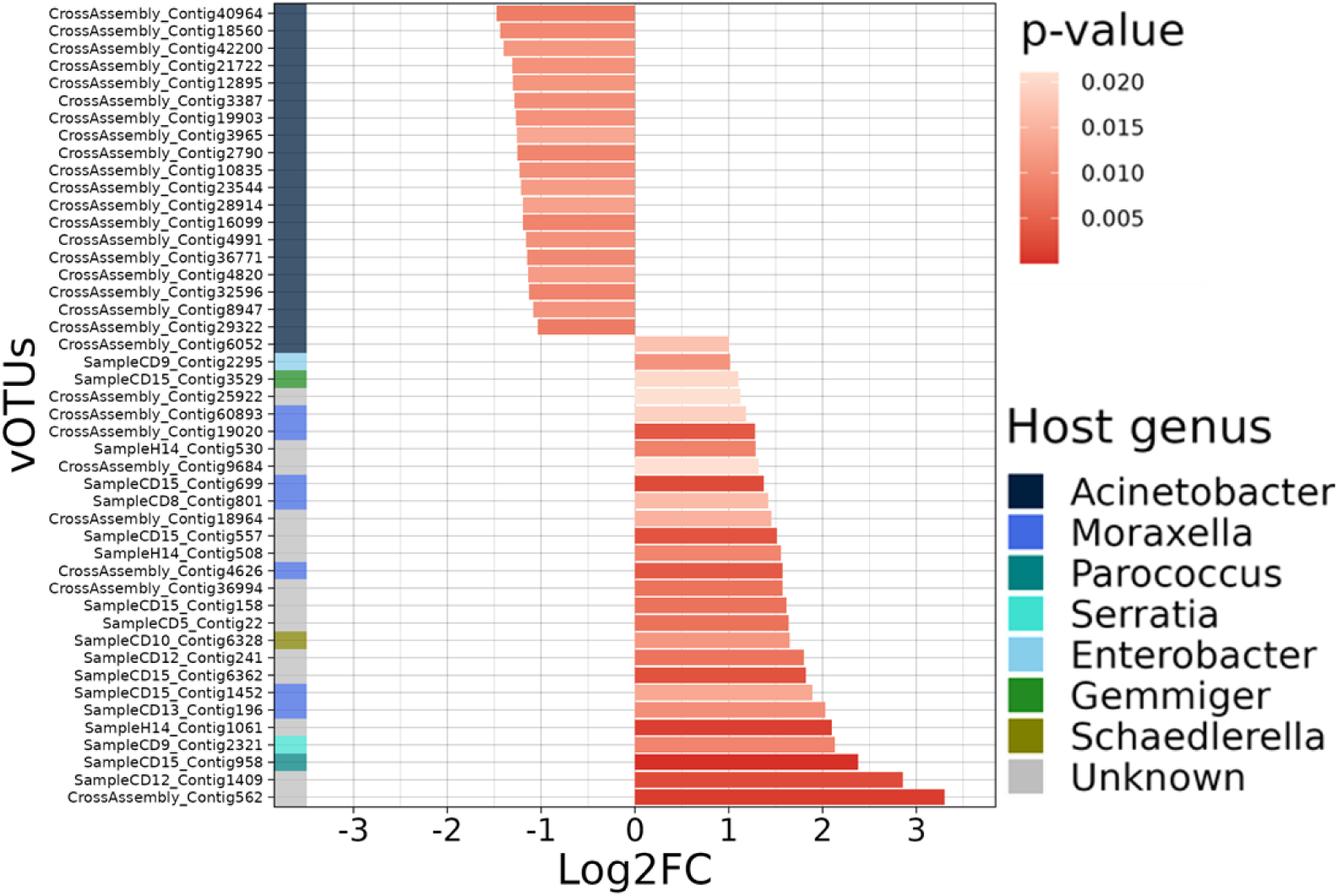
Differentially abundant blood vOTUs. Differential analysis was performed using Maaslin2. Only vOTUs with an adjusted p-value < 0.25 and Log2FC > 1 or < −1 were considered significant and are displayed.

To conclude, while a variety of phages are more abundant in the blood of CD patients, only the phages infecting Acinetobacter are found to be lower in numbers.

## Discussion

The viral component of the human blood remains largely unexplored, despite its vast potential as a marker of disease. This has been highlighted by recent studies showing that circulating phage DNA can be used to precisely identify bacterial pathogens causing sepsis^17^ or that the presence of specific phages in hemocultures could help detect pathogens^39^. In this study, we developed adequate methods to uncover a diverse phage community in the human blood. Most of the blood phages were predicted to infect Pseudomonadota bacteria, mirroring blood bacteria, dominated by the same phylum^1–3^. Recent microbiota studies have shown that the phage composition usually reflects the bacterial composition, as both populations co-exist over long periods of time^40^. Our observations follow this trend, which explains the dominance of phages infecting this phylum in this ecosystem. We additionally report the first quantification of viral particles in human blood, estimated at 10^5^ viral particles per mL of blood. Since the number of bacterial genomes predicted by 16S sequencing is around 10^6^ per mL^4,41^, the hypothetical phage:bacteria ratio of 1:10 in the blood matches the one reported in the human gut^42^, suggesting a similar organization.

However, despite the unexpected diversity and abundance of phages in the human blood we uncover, key questions remain about their impact on this ecosystem. With our virome sequencing approach, we are unable to distinguish infectious from inactive phage particles. Phage predation of blood bacteria is therefore difficult to assess with our data. The blood environment is known to affect phage-bacteria interactions and to reduce phage infectivity in some cases^43,44^, though large-scale studies are lacking to confirm these observations. This suggests that phages would not be able to replicate in this environment. The fact that bacteria isolated from the blood are in L-form or dormant metabolic states^2,4^ is another hindrance limiting phage replication in the blood. The presence of a blood phage community is therefore likely to have a limited effect on blood bacteria. Yet, such phages could have a direct impact on the human host, as exemplified through increasing examples of interactions between phages and the immune system^45–47^, including in the case of CD where ileal viromes of patients were found to exacerbate inflammation *in vivo*^48^. A phage infecting *Ralstonia* bacteria (Pseudomonadota phylum) was recently isolated from the blood and was shown to be persistent in this environment^49^, which is in line with our results. Remarkably, this phage codes for proteins which are capable to be expressed in eukaryotic cells and transported into the nucleus, pointing out the possible interactions between blood phages and the human host. We also observed significant variations in the blood virome composition between CD and healthy individuals, which could reflect interactions between blood viruses and the immune system, given that CD strongly affects the immune response. More generally, in tandem with eukaryotic viruses such as Anelloviruses, phages could play a role in the regulation of the immune system, as observed in the gut microbiota^50–52^. Altogether, much like for bacteria, the presence of phages in the blood could have important implications for human health, but remains to be better understood.

Another central interrogation is about the origin of the blood phages. Given that phage half-life in blood has been estimated to few hours in pharmacokinetics studies^53^, and with phage replication expected to be negligible in this environment, phage input must be constant and quantitatively high to match our observations. Prophage induction is likely not strongly contributing to the presence of phage particles in the blood, as we estimate that temperate phages do not constitute the majority of the community. Additionally, bacterial metabolism is also expected to be minimal in the blood environment and would prevent virion production. Instead, we suggest that the major contributor would be translocation from different body-sites. Translocation of phages across intestinal epithelial cells has been demonstrated *in vitro*^16^, and these findings are consolidated by the results of our parallel paper, demonstrating phage translocation across intestinal and endothelial cell layers, as well as *ex vivo* across murine intestinal tissues ^30^. This provides a potential mechanism for a gut origin of blood viruses. In the present report, we found that the gut virome is a notable and significant source of blood phages, with around 20% of blood phages homologous to gut phages. Likewise, studies focusing on the bacteria of the human blood found that the gut microbiota is an important source of blood bacteria, with bacterial translocation as an underlying mechanism^1,3,5^. Notably, only 3% of blood viruses were found in paired fecal samples. A tentative explanation could be that the fecal microbiota is not representative of the microbiota found at all sites of the GIT^54^, while phage translocation may happen throughout its different sections, according to their different permeability. In particular, the small intestine is known for its high prevalence of Pseudomonadota bacteria^54,55^, which corresponds to the most frequent host phylum of blood phages. Therefore, its higher permeability, compared with the large intestine^56^, could explain the prevalence of phages infecting this phylum in the blood, and their scarceness in faecal samples. While some of the gut viral databases include other types of samples, fecal samples largely dominate and the inclusion of more viruses from other sections of the GIT would probably improve our search for the origin of blood phages. Nonetheless, the origin of a large part of blood viruses remains unknown. We have identified a limited contribution from the oral virome, which may become more substantial as oral virus databases expand to match the richness of intestinal ones (around 170,000 viral species in our combination of gut databases compared to less than 50,000 in the Oral Virus Database), thereby enhancing our understanding of their connection to this ecosystem.

Other human-associated microbiota could also be explored to better understand the origin of blood phages. The skin hosts a well-known microbial community of yeasts, bacteria and phages^57^. However, the bulk of the organisms identified in the negative control samples was constituted of members of the skin microbiota such as *Malassezia* yeasts, *Propionibacterium* or other Actinomycetota bacteria, as well as their viruses. To ensure the reliability of our study, all these organisms were removed, which limits our ability to then detect skin viruses in our analysis. Nevertheless, the likelihood of viruses translocating through the skin appears to be low due to its stratified layers, including compact keratinocytes, and relatively low vascularity. Another more promising candidate could be the lung microbiota, which possesses a thin epithelium that facilitates gas exchanges and which could allow virus translocation. Lung epithelial cells were actually found to have the highest phage uptake in a study comparing seven different cell types^15^, highlighting the permeability of the lung epithelium and making the lungs a likely source of blood phages. Unfortunately, no database of lung viruses exists to date. The characterization of the viruses of this environment is ongoing^58^, which should enable to investigate its promising relation with the blood virome in the near future.

CD has been extensively studied in relation to the gut microbiota. Despite the lack of a specific marker of CD, significant changes in the gut microbiota of CD patients have been uncovered^20–22^. While earlier studies suggested similar changes in the gut virome^23–25^, recent analysis shows no significant differences between CD patients and healthy individuals^26^. Despite noting a decrease in viral alpha diversity among CD patients, which parallels the reduced bacterial diversity in this population, our observations also did not reveal significant disparities in the overall community composition. These findings therefore support the idea that the gut virome cannot be a marker of CD. The challenge most likely stems from the significant complexity and interindividual variability of this viral community^42^, making it difficult to identify a unified signal of disease.

Conversely, microbial communities outside of the gut tend to exhibit less complexity, which could facilitate the identification of a CD-specific signal. We thus chose the blood virome, which contains few viruses and is still unexplored. Our findings reveal for the first time an association between blood virome and a non-infectious disease. CD causes a myriad of changes in the immune status of the patients, as reflected by increases in inflammatory markers routinely measured in the blood such as the C-reactive protein. How these alterations of the immune system can in turn affect the phage population remain difficult to predict. Besides, CD is also known to undermine the gut barrier integrity and thus its permeability, which could lead to changes in phage transfer to the blood. In a parallel article, we show that viruses are more frequently found in both fecal and blood samples of the same individual for CD patients ^30^. This confirms the increased transfer of gut phages into the blood in CD, potentially underpinning the changes in the blood virome observed in CD patients. Other studies have reported associations between blood bacterial communities and diseases affecting the immune system, such as IgA nephropathy^41^ or acute pancreatitis^3^. Our work further reinforces the nascent link between blood microbes and disease by integrating the viral component.

We were able to precisely identify a group of differentially abundant phages in CD patients. Strikingly, all the underrepresented phages had *Acinetobacter* as predicted host. This constitutes a clear signal of the effect of CD on blood viral communities and establishes a novel signature of disease. *Acinetobacter* has been reported to be one of the most represented bacterial genera found in the blood in multiple studies, with relative abundances from 3 to 10%^3–5^. Changes in the population of phages infecting this genus could therefore have strong implications on this ecosystem. If the virome reflects the bacteriome, one could expect that the abundance of the bacteria is likewise reduced in the blood of CD patients. Nevertheless, current knowledge of blood bacteria is limited and further work, such as sequencing both bacterial and viral DNA of blood samples, is required to better understand of the significance of this promising observation.

To conclude, our results uncover unsuspected phage diversity in the blood virome, advancing our understanding of this environment. This led to the identification of a different blood viral community in CD patients, which could be used as a signature of CD and improves our comprehension of this complex disease. Altogether, our findings pave the way for further exploration of the virome and microbiome of the blood in context of other disorders.

## Material and Methods

### Cohort

CD patients and healthy individuals were recruited through the Gastroenterology Department of the Saint-Antoine Hospital (Paris, France). All patients provided their informed consent, and the study was approved by the corresponding ethics committee (Comité de Protection des Personnes Ile-de-France IV, IRB 00003835 Suivithèque study; registration number 2012/05NICB). Our cohort contained 15 CD patients and 14 healthy controls. None of the individuals underwent intestinal surgery, and none took any antibiotic treatment in the last 3 months before sample collection. The median age of the cohort was of 38 ± 11 years. Detailed information about all the individuals of the cohort can be found in **Table S1**.

### Sample collection

Comparing serum and plasma for the retrieval of phages (samples were spiked with T4, lambda and M13 phages) showed no major difference. Therefore, plasma was chosen for its easier downstream processing. Blood was collected in blood collection tubes designed for plasma collection (Dutscher ref. 369032), and immediately put on ice upon drawing. The samples were then centrifuged at 2,000 g for 10 min to separate the plasma. Plasma was then filtered through a 0.45 µm cellulose acetate filter, and stored at −20°C.

Fecal samples were collected directly in RNAlater (Sigma ref. R0901) and homogenized by vortexing using glass beads (Sigma ref. Z265942). After a centrifugation at 20,000 g for 15 min at 4°C, the supernatant was removed, and the pellet was directly stored at −80°C. RNAlater contains a high concentration of ammonium acetate, which precipitates phages particles. Phages are therefore contained in the pellet after centrifugation. Blood and fecal samples from the same individual were collected on the same day.

For negative controls, a blood collection tube and a low-binding Eppendorf tube (ref. EP0030108426) containing sterile, double-distilled water were used.

### Viral DNA preparation and sequencing

To extract the viral DNA from blood and fecal samples, we adapted a virome protocol from Billaud *et al*^59^. Fecal samples (0.2 g) were thawed on ice, resuspended in 7 mL of Tris 10 mM pH 7.5. Plasma samples (3 mL) were thawed on ice. After this step, all samples were processed similarly. First, the samples were centrifuged at 4,750 g for 10 min at 4°C to remove most bacterial cells. The supernatant was then filtered through a 0.45 µm PES membrane filter to remove remaining bacteria. At this step, samples were spiked with known quantities of phages SPP1 (dsDNA) and M13 (ssDNA) (see “Estimation of the number of viral particles in samples by spiking” section). The negative control samples were not spiked. A benzonase-nuclease treatment (250 U, Sigma ref. E1014) was then performed for 2h at 37°C. In order to concentrate viruses, samples were mixed with 0.5 M NaCl and 10% weight/volume PEG-8000, followed by overnight incubation at 4°C. Samples were then centrifuged at 4,750 g for 30 min at 4°C. The precipitate was collected and re-suspended in 400 µL Tris 10 mM pH 7.5, and shaken gently with an equal volume of chloroform in order to kill remaining bacteria and reduce vesicle contamination. After transferring in Maxtract gel tubes (Qiagen ref. 129056), the samples were centrifuged at 16,000g for 5 min at 4°C, after which the aqueous phase was extracted. To remove non-encaspidated DNA, samples were treated with TURBO DNAse I (8 U, ThermoFisher ref. AM2238,) and RNAseI (20 U, ThermoFisher ref. EN0601). DNAse was then inactivated using EDTA at a final concentration of 20 mM. To break open the capsids, proteinase K was added (40 µg) with 20 µL of SDS 10% before incubation at 56°C for 3 h. The samples were transferred into Maxtract gel tubes containing an equal volume of Phenol-Chloroform-Isoamyl alcohol (25:24:1, ThermoFisher). After vigorous vortexing, samples were centrifugated at 16,000 g for 5 min at 4°C. The aqueous phase was extracted and transferred to a new Maxtract gel tube. To remove phenol traces, chloroform/isoamyl alcohol (24:1, ThermoFisher) was added, followed by another identical centrifugation. The aqueous phase was extracted and DNA precipitated using sodium acetate (0.3 M), ethanol (2.5 volumes) and glycogen (20 µg/mL final concentration), then incubated 1 h at −20°C. The samples were then centrifuged at 16 000 g, 30 min, 4°C. The pellet was washed twice with 0.5 mL cold 75% ethanol. After ethanol removal, the dry pellet was resuspended in Tris 10 mM pH 8 and stored at −20°C.

Libraries were then prepared using the xGen kit, which provides a step to convert ssDNA to dsDNA with minimal biais^60^. The resulting libraries were then sequenced on an Illumina apparatus (paired-ended 150 bp reads). The sequencing generated 9.2 million (**±** 6 million) pairs of reads (**Table S2**).

### Sequencing QC and human read removal

To remove low quality reads and the “adaptase” tail characteristic of the xGen kit, raw sequencing reads were processed using fastp version 0.23.1^61^ with the following settings: *--trim_front1 2--trim_tail1 2--trim_front2 15--trim_tail2 2 -r -W 4 -M 20 -u 30 -e 20 −l 100*.

The reads of the human host were removed by mapping the cleaned reads against the human genome (GRCh38 version) using bowtie2 version 2.4.1^62^ with the *--very-sensitive* option. After this step, four samples had less than 100,000 pairs of reads remaining (H7_F, H10_F, H13_B, CD7_B) and were thus removed from subsequent analysis.

### Assembly

The assembly was performed using the cleaned, non-human reads both per individual (using blood and fecal samples together), and with the reads from all the samples of all the individuals altogether (cross-assembly). In both cases, both paired-end and unpaired reads were used, and all the reads were deduplicated prior to the assembly using the dedupe function of bbmap version 38.86^63^.

The assembly was then performed using SPAdes version 3.14.0^64^ with default parameters, which slightly outperformed metaspades and metaviralspades in number and average size of contigs. Quality of the assemblies was assessed using QUAST version 5.2.0^65^. Contigs < 2 kb, which correspond to the minimal size of DNA viral genomes, were removed.

### Delimitation of OTUs by clustering

Contigs were clustered to delimit OTUs using an approach adapted from Shah et al^35^. First, pairwise alignment of all the contigs was performed using BLAT version 36^66^. The contigs having a total alignment length against themselves > 110% were considered chimeras and removed. For the remaining contigs, clustering was performed at the 95% average nucleotide identity (ANI), corresponding to species boundaries, using scripts described in Shah et al^35^.

### vOTU detection

Two approaches were combined to detect vOTUs: dedicated tools and homology to virus databases. For the tools, VirSorter2 version 2.2.3^67^ was used with the following parameters to increase stringency: *--include-groups dsDNAphage,NCLDV,RNA,ssDNA,lavidaviridae--exclude-lt2gene--min-length 2000--viral-gene-required--hallmark-required--min-score 0.9--high-confidence-only*. We included in our viral dataset all the OTUs predicted to be viral. VIBRANT version 1.2.1^68^ was also used, and all OTUs predicted as viral were included. CheckV version 0.8.1^28^ was also run on all of the OTUs. Only the OTUs with a viral quality above “medium” were included. The comparison between the outputs of these three tools is represented in **Fig. S1**.

For the databases, the OTUs were compared to the COPSAC^35^, the GVD^31^, the MGV^32^, the GPD^33^, RefSeq Virus^69^ (12/07/2023 release) and the CHVD^34^ databases using BLAT version 36^66^. OTUs having > 90% ANI homology to at least one database were considered viral. The comparison between the outputs of the five gut databases is displayed in **Fig. S2**. Finally, the vOTUs of both methods were combined to form a non-redundant list of 17,454 viral vOTUs (**Fig. S3**).

### vOTU annotation

The vOTUs were annotated using a modified version of the RimeTOOLS2 Pipeline, from Rime Bioinformatics. Coding sequences (CDS) and tRNAs were predicted using the following tools: prodigal 2.6.3^70^ (option-meta), phanotate 1.5.0^71^ (default parameters), glimmer 3.02^72^ (options long-orfs: -n -t 1.1.15; -o 50 -g 110 -t 30), genemarks 1.14^73^ (options -m heu_11.mod, -r) and tRNA-scan-SE 2.0.7^74^ (option-B). tRNAs were detected first and used to mask the genome sequence, before a 2-tool consensus prediction of CDS using prodigal, phanotate, glimmer, and genemarks. Only CDS called by at least two independent tools were considered, and the longest CDS was prioritized in case of conflict. CDS were subsequently used to query gene and protein databases (CARD 05/2023^75^, Resfinder 04/2023^76^, VFDB 05/2023^77^, Phantome 03/2021, Swissprot 05/2023^78^, NCBI Virus 15/2023, Vog 05/2020, Pvog 03/2021^79^, Refseq 05/2023^69^, DefenseFinder 05/2023^80^, and PHROGs 05/2023^81^) using HMMER 3.3.2^82^ (E-value < 0.00001, Coverage > 0.7), HHsuite 3.3.2^83^ (E-value < 0.00001, Coverage > 0.7, Probability > 90%, Score > 30), and Blast+ 2.9.0^84^ (E-value < 0.00001, Identity > 0.7, Coverage > 0.7). Consensual results with high match scores were kept for the final functional annotation, and prioritized depending on the database of origin (PHROGs > Resfinder, VFDB, DefenseFinder, CARD, Swissprot > Phantome, pVOG, VOG > Refseq).

### vOTU classification

To classify the vOTUs, viral proteins (see “vOTU annotation” section) were combined with those of the INPHARED database^85^ (02/07/2023 release), and used as input for clustering using vContact2 version 0.9.19^86^ using default parameters. Then, graphanalyzer version 1.5.1^87^ was used to assign a classification from the network generated by vContact2. A classification was obtained for 2,217 vOTUs (13% of vOTUs).

### vOTU lifestyle prediction

The prediction of the lifestyle for the vOTUs was done by searching for integrases in the annotated viral genomes. Only high-quality or complete genomes, as assessed by CheckV, were considered for this analysis. vOTUs classified as Microviridae were also excluded, as their lifestyle cannot be predicted by the presence of integrases^88^. Genes were considered as integrases when either: “integrase” was present in the gene product name, “int” was the gene name, or when both “recombinase” and either “serine” or “tyrosine” are present in the gene product name. vOTUs with at least one integrase were considered as having a temperate lifestyle.

### vOTU host prediction

Host prediction for vOTUs was performed using iPHoP version 1.2.0^89^ with the associated database (09/2021 release) using default parameters. For both “genome” (host-based tools only) and “genus” (host- and phage-based tools), the best hit for each vOTU was kept. When a vOTU had a hit for both modes, the “genus” prediction was favored. A host was predicted for 13,856 vOTUs (79% of vOTUs).

### Non-viral OTUs classification

All OTUs, including the non-viral OTUs, were classified using CAT version 5.2.3^90^ using default parameters.

### Identification of eukaryotic viruses

All OTUs were compared to the RefSeq Virus database using BLAT (see “vOTU identification” section). The OTUs homologous to RefSeq Virus were hand-curated to removed viruses of bacteria and archaea in order to identify the eukaryotic viruses.

### Relative abundances calculation

Cleaned, non-human reads were mapped using bwa-mem2 version 2.2.1^91^ against all the OTUs, using default parameters. The resulting SAM file was converted to BAM and sorted by read name using samtools version 1.9^92^, before filtering out the low-quality alignments using msamtools filter version 1.1.3. Alignments > 80 bp with > 95% identity over > 80% sequence length were selected, and only the best alignment (or alignments in the case of multiple alignments with the best score) for each read were kept. The alignments were transformed into counts using msamtools profile, with the *multi=proportional* option so that reads mapping to different OTUs were allocated to the OTUs relatively to their abundances. The coverage and depth of each OTU for each sample was determined using msamtools coverage. OTUs for which sequence coverage was either smaller than 50% of the OTU size or the average sequencing depth was smaller than 1 in a given sample were discarded by setting their count to 0 in this sample. The OTUs present in the negative control samples were set to 0 counts in all the other samples to remove contaminants. Given the disparity in number of viral counts for blood and fecal samples and their separate analysis, rarefaction was performed separately for each. For fecal samples, counts were rarefied at the lowest number of viral counts (267,514 counts). For blood samples, counts were rarefied at 7,653 viral counts (three samples were excluded by this rarefaction). Counts were then normalized by vOTU length and transformed into relative abundances to allow for comparison across vOTUs and samples. This resulted in the main abundance table used for the analysis.

For the analysis of negative control samples and of non-viral OTUs, a similar approach was performed. After the coverage criteria were applied, the entire count table was directly normalized by OTU length and transformed in relative abundances before analysis.

### Ecological analysis

The relative abundance table was analysed in tandem with a taxonomy table, containing all the information gathered about each vOTU (length, quality, predicted host, annotation,…) with the R package phyloseq version 1.46.0^93^. The R package vegan version 2.6.4^94^ was used for the Adonis2 multivariate analysis.

### Estimation of the number of viral particles in samples by spiking

In order to estimate the number of viral particles in the samples, we employed a spiking approach using two phages: a dsDNA phage, SPP1 (host *Bacillus subtilis*) and a ssDNA phage, M13 (host *Escherichia coli*). In fecal samples, a concentration of 10^6^ plaque-forming units (PFU) per mL of both SPP1 and M13 was used for all samples. In blood samples, two concentrations were used: 10^6^ PFU/mL for both SPP1 and M13 for samples H1 to H5 and CD1 to CD5, and 10^4^ PFU/mL for the other samples. The vOTUs corresponding to SPP1 and M13 were identified by homology search against the reference genomes of these phages, using vsearch version 2.18.0^95^. Then, for each sample, the relative abundance of both SPP1 and M13 was calculated using the corresponding concentration spiked in the sample, in order to generate an estimation of the number of viral particles contained in the rest of the sample (contaminant and non-viral OTUs were removed before this step). Then, the values obtained for SPP1 and M13 were averaged. This number was finally normalized by the weight of the fecal sample (0.2 g for all samples) to generate viral particles per g of feces estimates for fecal samples, or by the volume of blood used (3 mL for all samples) to generate viral particles per mL of blood for plasma samples.

### Determination of the origin of the blood vOTUs

In order to determine the origin of the vOTUs, the comparison of the OTUs against gut viral databases (COPSAC, CHVD, GVD, GPD) was used (see “vOTU detection” section). With a threshold of > 90% ANI, blood vOTUs having homology against at least one these databases were considered as having a gut origin. The Oral Virus Database^37^ was used in the same manner in order to find blood vOTUs having an oral origin.

### Differential analysis

To identify vOTUs responsible for the different blood viral communities in CD patients and healthy individuals, differential analysis was conducted using the Maaslin2 R package^96^. The minimum prevalence was of five samples, and no minimal abundance was set. Normalization was performed by total sum scaling, and the analysis method was LM (both are default parameters). The fixed effects were the disease status (CD/healthy) and the sex. The age, which was statistically lower in the control group (median age: 42 in the CD group and 30 in the healthy group, p=0.0027, one-way ANOVA), could not be added in the model. Only the vOTUs for which q-value < 0.25 and |Log2FC| > 1 were considered significant.

### Statistical analysis

All statistical analysis was performed using R version 4.3.2^97^. On the boxplot displayed in the figures, the middle line represents the median value, the bottom and top edge of the box represent respectively the lower and upper quartiles, and the whiskers extends from the edges of the box to the largest or smallest value no further than 1.5 times the inter-quartile range. Pairwise comparisons were performed with the Wilcoxon test.

## Acknowledgements

We thank Nicolas Dufour for valuable discussion and his suggestion to include the blood collection tube as a negative control. We thank Marius Bredon for his advice on the differential analysis. We thank Sophie Thenet, Véronique Carriere, Yanis Sbardella and Clara Douadi for valuable discussions. We thank Eugen Pfeifer for critical review of the manuscript. We also want to thank Rio for his contribution to the virome protocol. We are grateful to the INRAE MIGALE bioinformatics facility (MIGALE, INRAE, 2020. Migale bioinformatics Facility, doi: 10.15454/1.5572390655343293E12) for providing computing and storage resources. Finally, for the sequencing, we thank M. Monot, GM Haustant, L. Lemée Biomics Platform, C2RT, Institut Pasteur, Paris, France, supported by France Génomique (ANR-10-INBS-09-09) and IBISA. Some of the figures were created using BioRender.com.

## Data availability

The sequencing data have been deposited on ENA (PRJNA1126425). The vOTU annotations are available at https://doi.org/10.57745/FTIXS5.

## Code availability

The data and code used to produce all the figures can be accessed at https://github.com/QLamyBesnier/Human_Blood_Phageome_2024.

