## Supplementary figures and tables for "The human blood harbors a phageome which differs in Crohn’s disease"

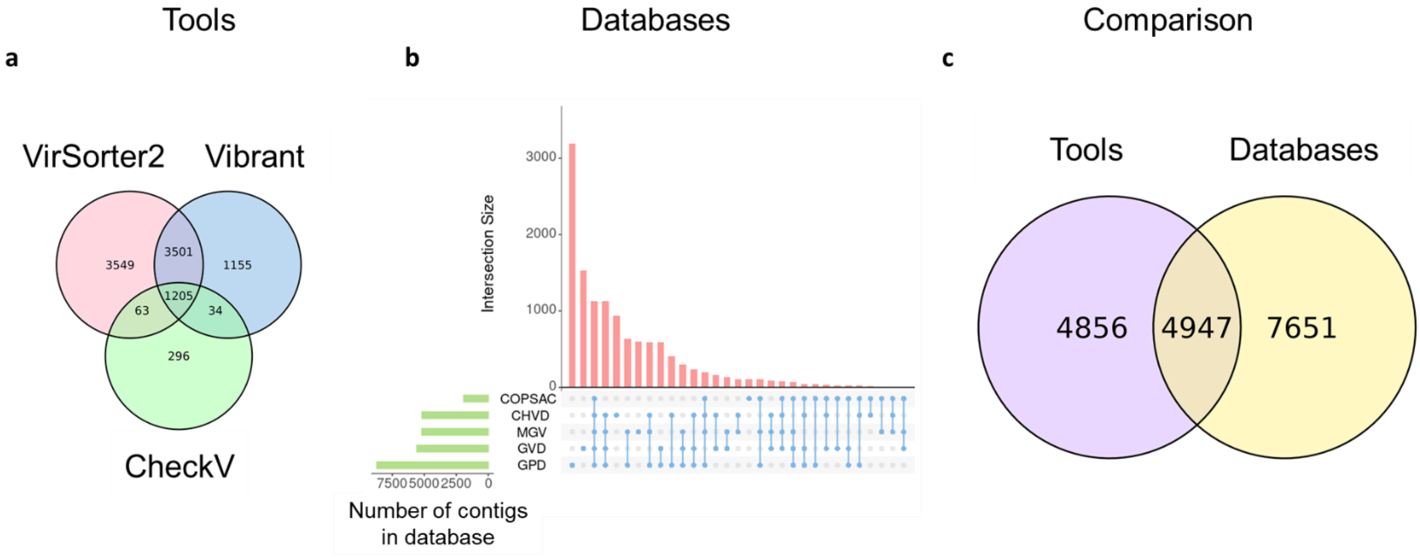


**Figure S1: Detection of vOTUs.**
**a.** Venn diagram of the OTUs detected as viral by VirSorter2, VIBRANT and CheckV. **b.** Upset plot of the OTUs detected as viral through homology with five different gut virus databases. **c.** Venn diagram of the OTUs detected as viral by the tools (left) and the databases (right).

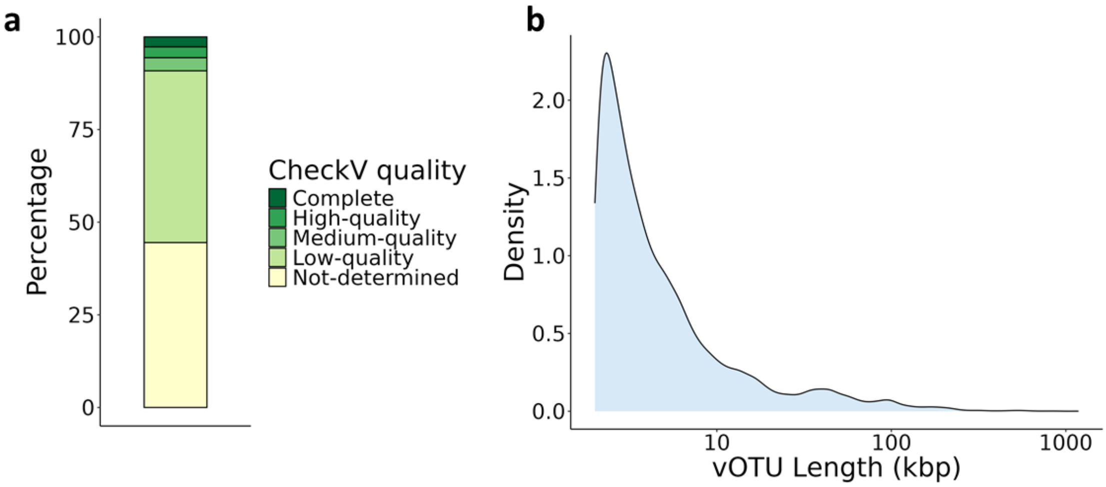


**Figure S2: Characteristics of vOTUs.**
**a.** Quality of the vOTUs as determined by CheckV. **b.** Distribution of the length of the vOTUs.

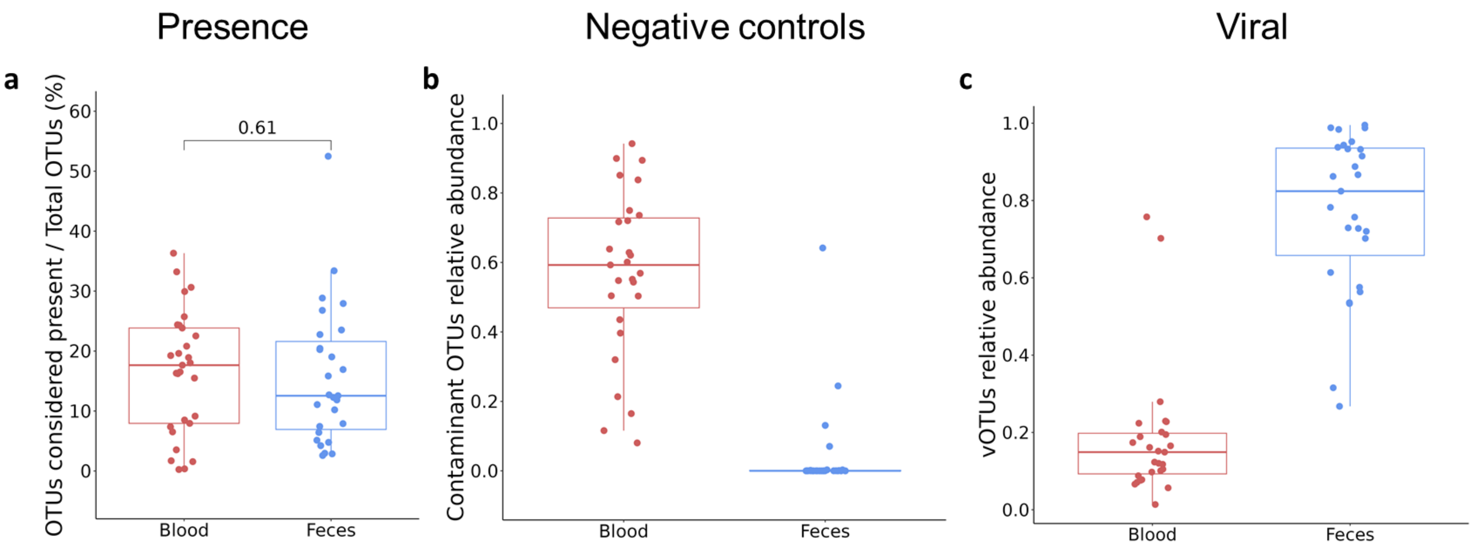


**Figure S3: Effects of the different filters used to select non-contaminant vOTUs.**
**a.** Proportion of OTUs that are considered "present" (depth > 1 and coverage > 50%, Methods) per sample, separated by sample type. **b.** Total relative abundance of OTUs that are considered contaminant (i.e. present in negative control samples) per sample, separated by sample type and disease status (orange = CD patients, green = healthy individuals). **c.** After removal of contaminant OTUs (panel b), relative abundance of the OTUs that are considered viral, separated by sample type and disease status (orange = CD patients, green = healthy individuals).

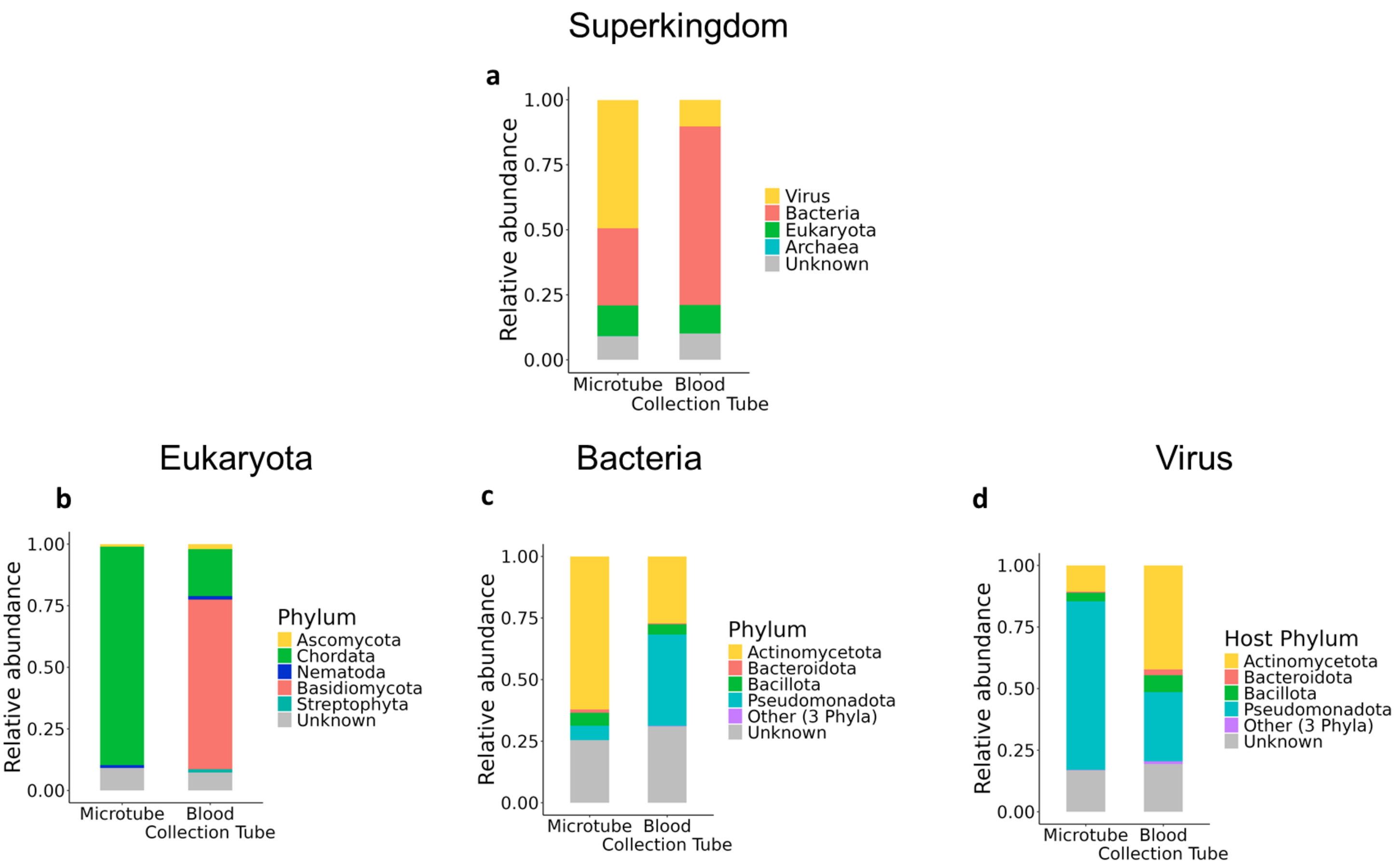


**Figure S4: Composition of the negative controls.**

**a.** Composition of both negative control samples, at the superkingdom level. **b.** Composition of the Eukaryota fraction of both negative control samples, at the phylum level. **c.** Composition of the Bacteria fraction of both negative control samples, at the phylum level. **d.** Composition of the Viral fraction of both negative control samples, classed by the predicted host phylum level.

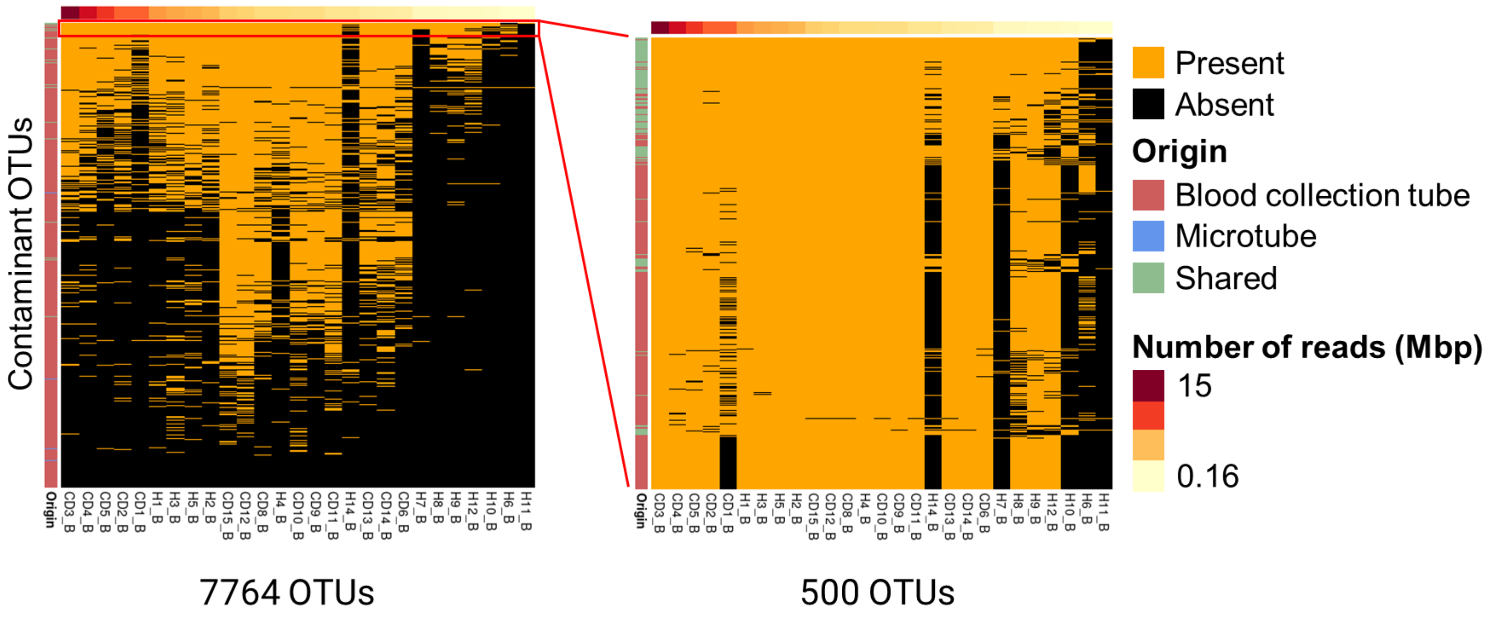


**Figure S5: Contamination across samples.**
Presence/absence heatmap of the contaminant OTUs (present in negative control samples) across the blood samples. The OTUs are organised top to bottom by decreasing sharedness, and the samples are organised left to right by decreasing number of non-human, cleaned reads. A zoom on the 500 most shared OTUs is displayed on the right panel. The origin (i.e. the sample(s) in which the OTU was found to be present) of each vOTU is displayed on the left of the heatmap.

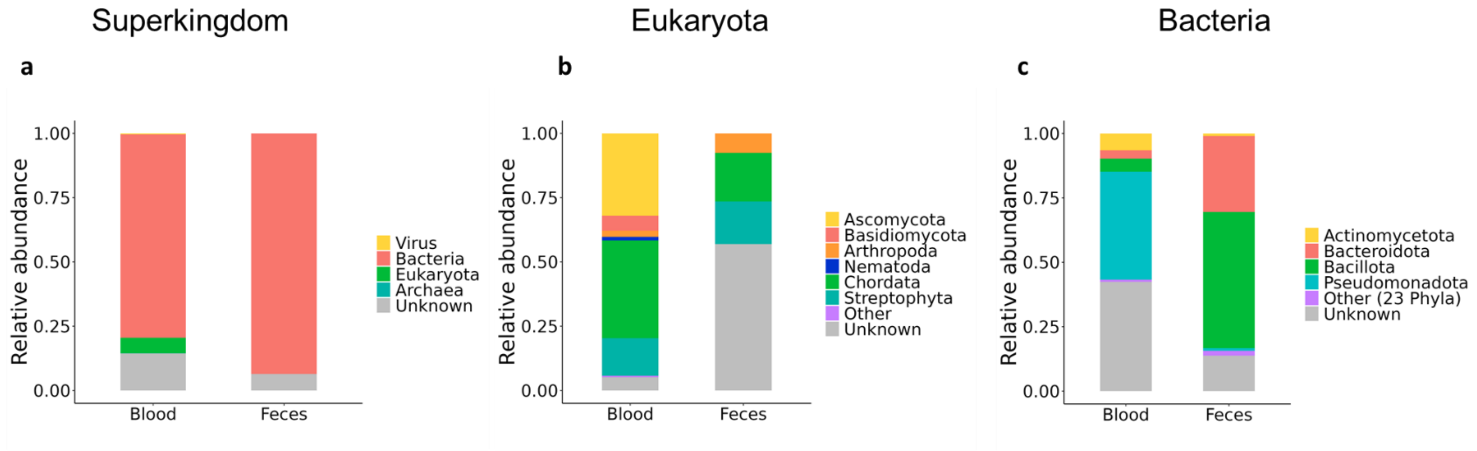


**Figure S6: Composition of the non-viral fraction of the samples.**
**a.** Average composition of the non-viral fraction of the samples at the superkingdom level, separated by sample type. **b.** Average compositon of the eukaryotic fraction of the samples at the phylum level, separated by sample type **c.** Average composition of the bacterial fraction of the samples at the phylum level, separated by sample type.

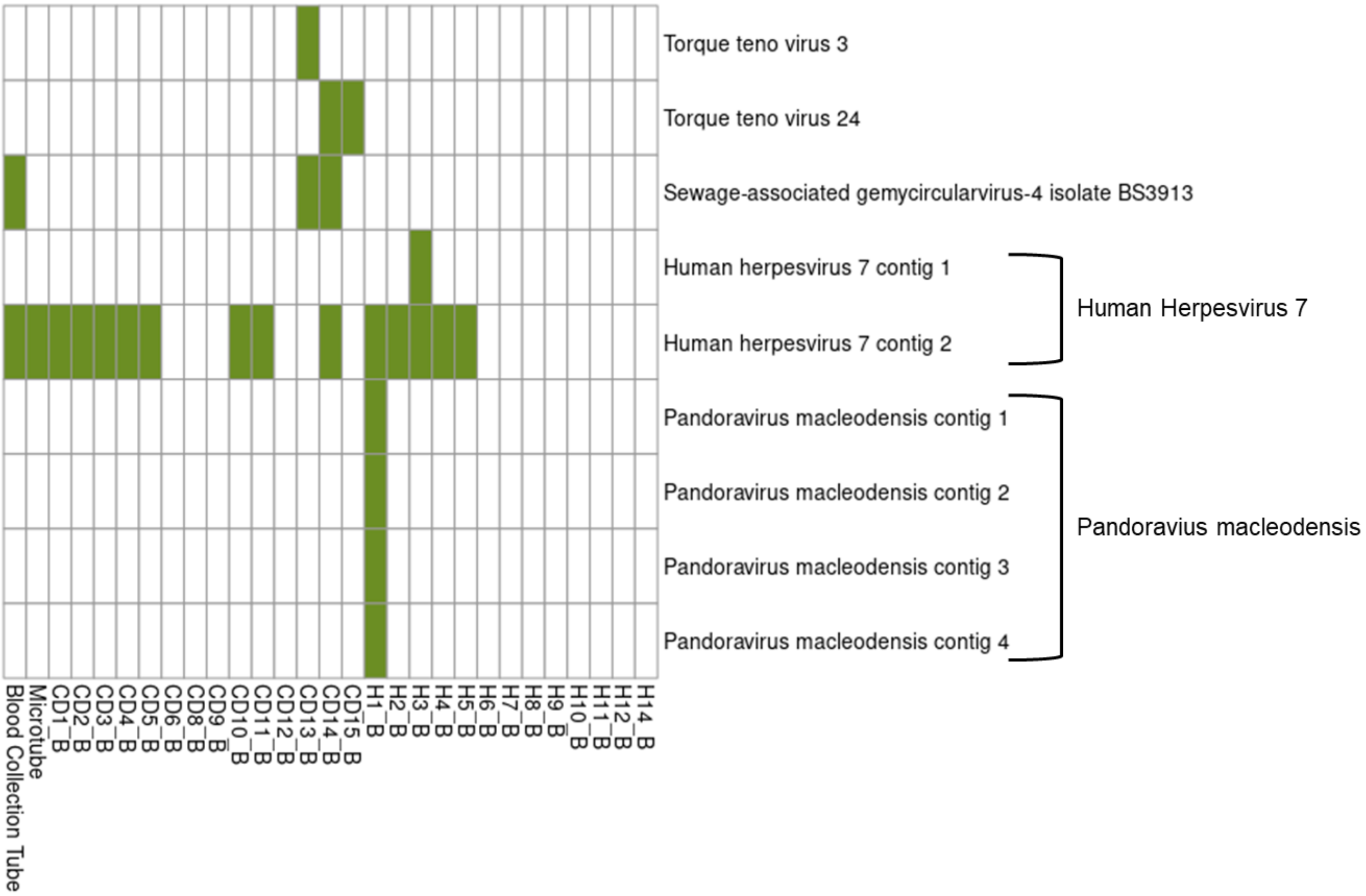


**Figure S7: Eukaryotic viruses in blood samples.**
Presence/Absence table of the eukaryotic viruses found in the negative controls and in the blood samples.

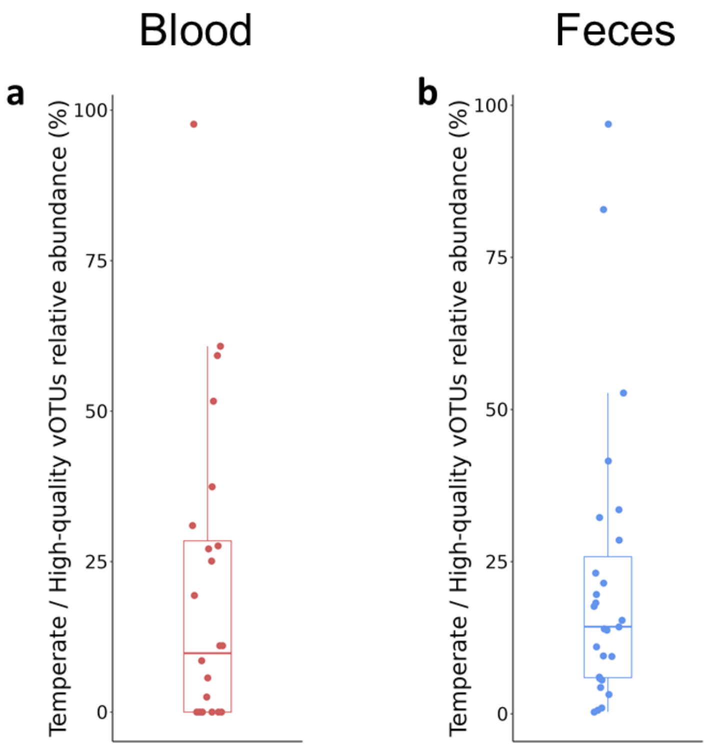


**Figure S8: Temperate phage content of blood and fecal samples.**
**a.** Relative abundance of temperate phages among all high quality vOTUs, per blood sample.
**b.** Relative abundance of temperate phages among all high quality vOTUs, per fecal sample.

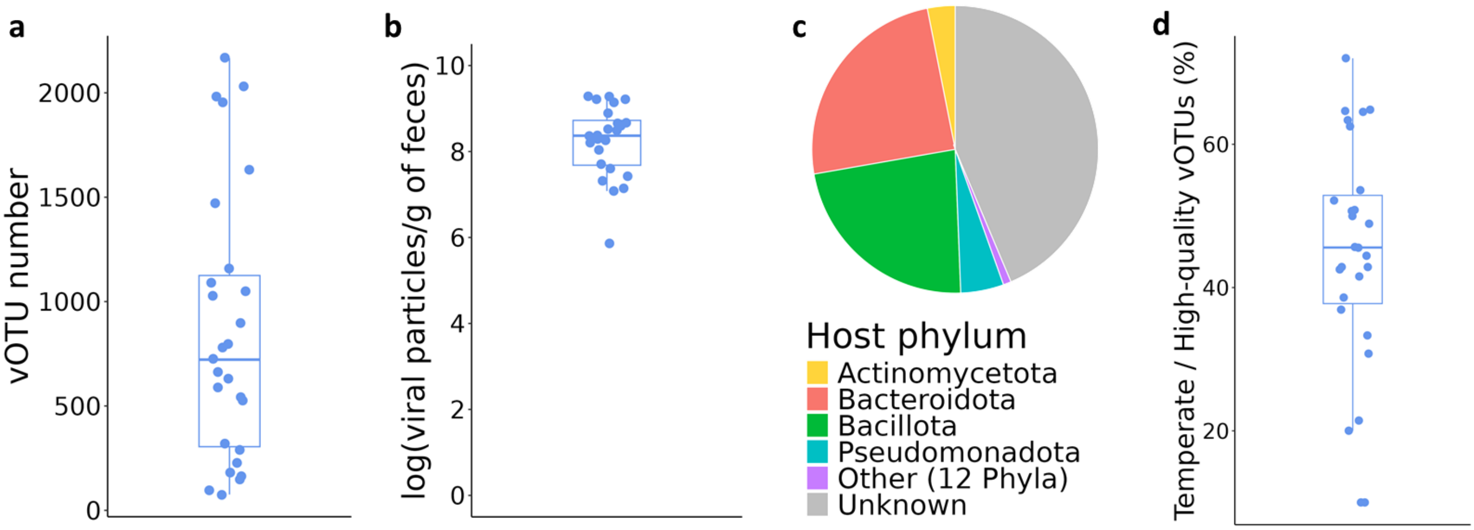


**Figure S9: Characteristics of the human fecal virome.**
**a.** Number of vOTUs per fecal sample. **b.** Estimated number of viral particles per g of feces and per fecal sample. **c.** Distribution of the predicted host phyla for all the vOTUs found in fecal samples. **d.** Percentage of vOTUs with a temperate lifestyle among the high-quality vOTUs. The temperate lifestyle was predicted by the presence of genes coding for an integrase, and only the high-quality genomes were considered (Methods).

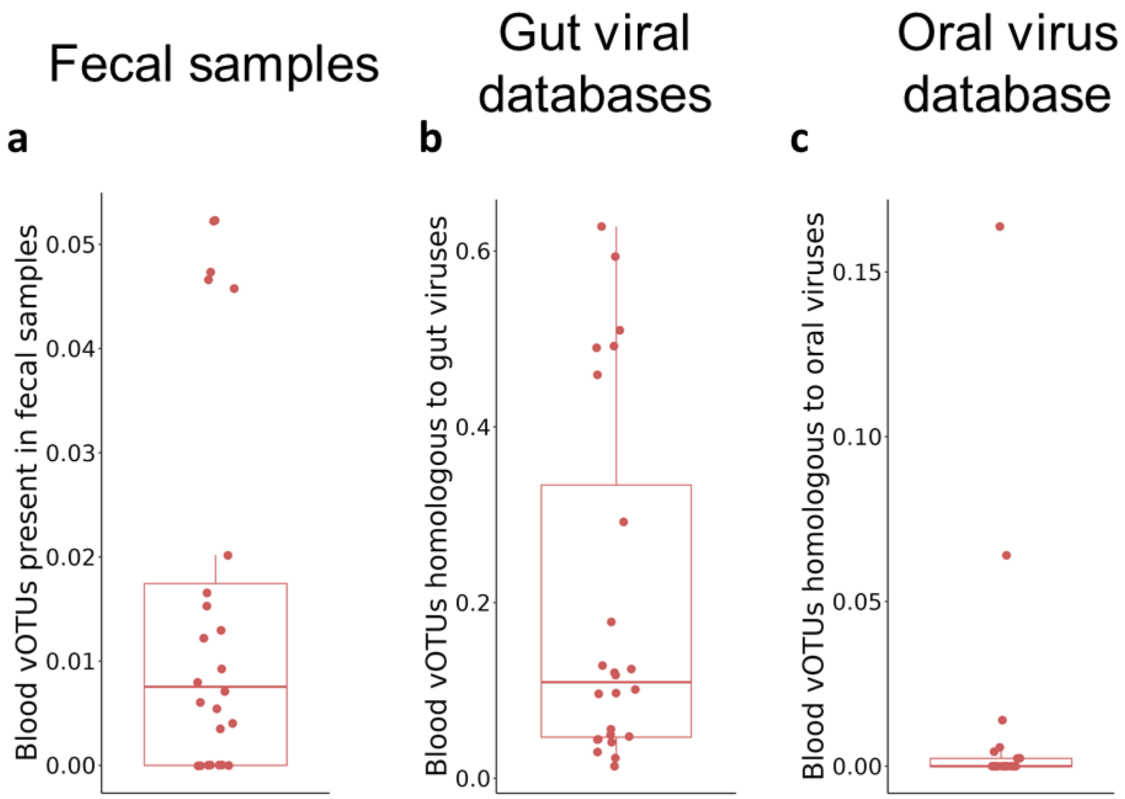


**Figure S10: Abundances for the blood vOTUs of different origins.**
**a.** Total relative abundance per blood sample of the vOTUs also found in fecal samples. **b.** Total relative abundance per blood sample of the vOTU homologous to viruses found in gut virus databases. **c.** Total relative abundance per blood sample of the vOTU homologous to viruses found in the Oral Virus Database.

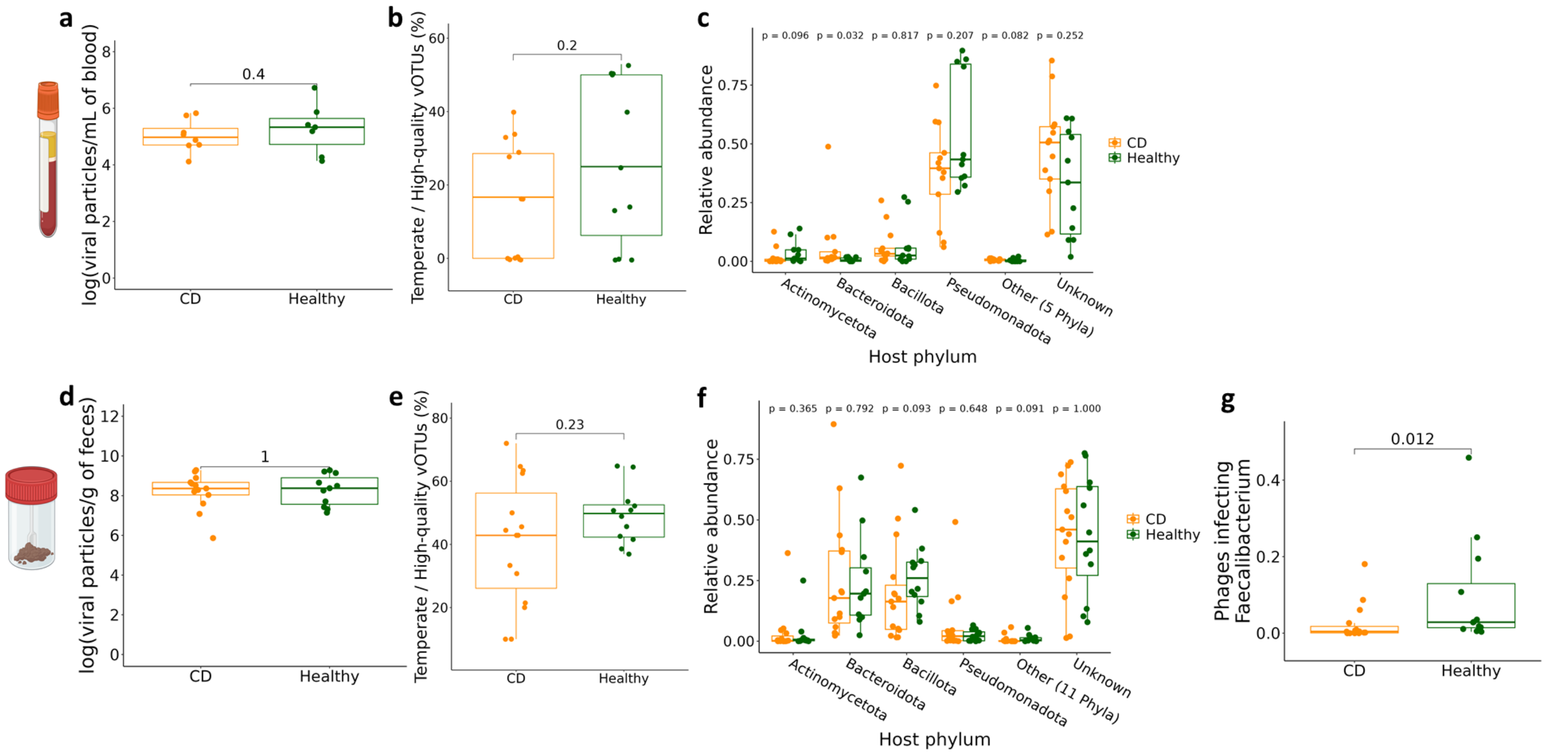


**Figure S11: Crohn's disease vs. healthy comparison for blood virome metrics.**

**a.** Estimation of the quantities of viral particles per mL of blood and per blood sample, separated by disease status. **b.** Percentage of vOTUs with a temperate lifestyle among the high-quality vOTUs per blood sample, separated by disease status. The temperate lifestyle was predicted by the presence of genes coding for an integrase, and only the high-quality vOTUs were considered (Methods). **c.** Total relative abundance for each predicted host phylum, per blood sample, separated by disease status. **d.** Estimation of the quantities of viral particles per g of feces and per fecal sample, separated by disease status. **e.** Percentage of vOTUs with a temperate lifestyle among the high-quality vOTUs per fecal sample, separated by disease status. The temperate lifestyle was predicted by the presence of genes coding for an integrase, and only the high-quality vOTUs were considered (Methods). **f.** Total relative abundance for each predicted host phylum, per fecal sample, separated by disease status. **g**. Total relative abundance for all phages predicted to infect the genus *Faecalibacterium*, per fecal sample, separated by disease status.

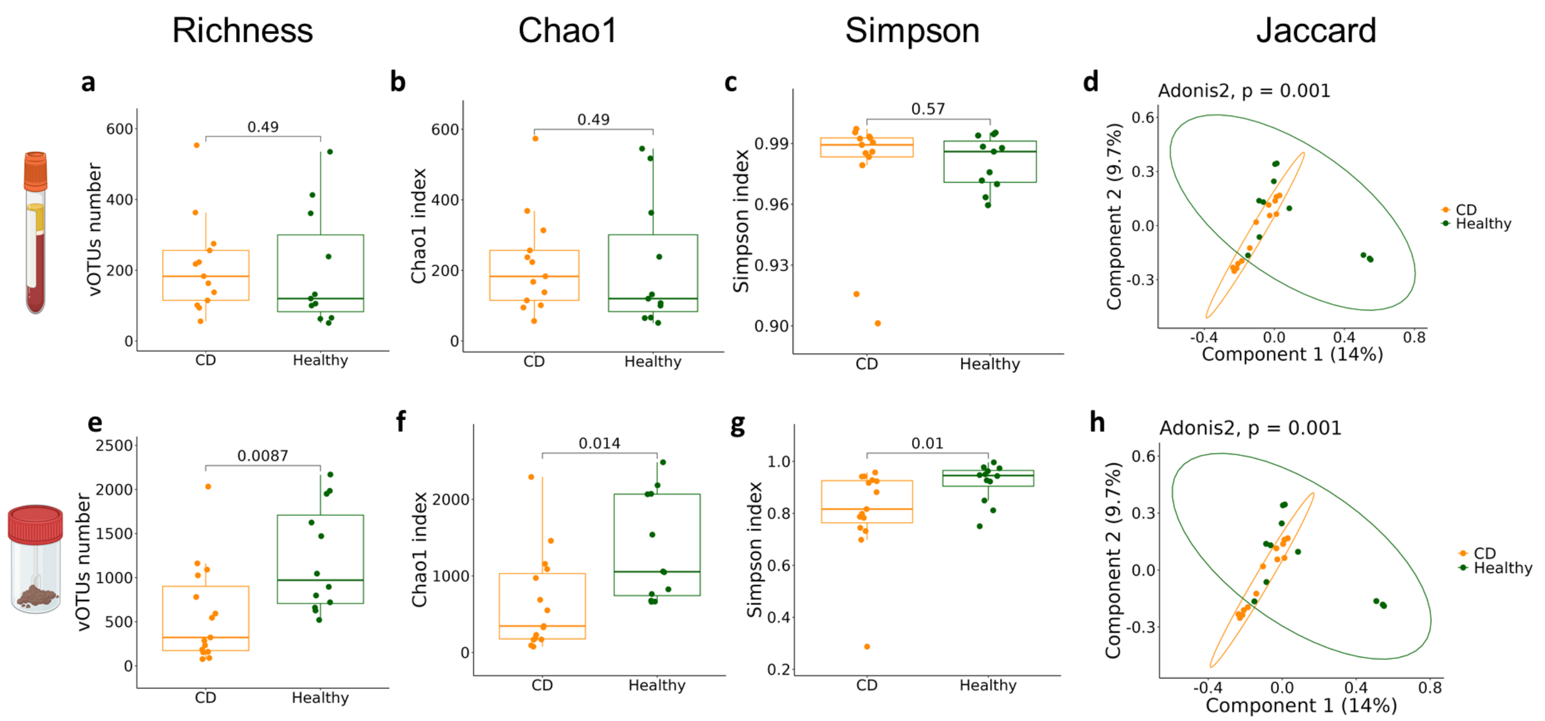


**Figure S12: Additional alpha and beta diversity measurements for the blood and fecal viromes.**

**a.** Number of vOTUs per blood sample, separated by disease status. **b.** Alpha diversity, measured by the Chao1 index for the blood samples, separated by disease status. **c.** Alpha diversity, measured by the Simpson index for the blood samples, separated by disease status. **d.** PCoA of the vOTU composition of the fecal samples, using Jaccard distances. The associated Adonis2 analysis revealed a difference between both groups (p=0.001). **e.** Number of vOTUs per fecal sample, separated by disease status. **f.** Alpha diversity, measured by the Chao1 index for the feces samples, separated by disease status. **g.** Alpha diversity, measured by the Simpson index for the fecal samples, separated by disease status. **h.** PCoA of the vOTU composition of the fecal samples, using Jaccard distances. The associated Adonis2 analysis revealed no significant difference between both groups (p=0.492).

### Supplementary Tables

| Individual | Sampling date | Sex | Age | Age at diagnosis | Localisation (Montreal) | Flare/Remission | C-reactive protein (mg/L) | Harvey-Bradshaw Index | smoking | Surgery | Antibiotics (last 3 months) | Undergoing treatment |
|---|---|---|---|---|---|---|---|---|---|---|---|---|
| CD1 | 29/02/2020 | Female | 53 | 37 | L2 | Remission | 1 | 2 | Past | No | No | Yes |
| CD2 | 2020 | Female | 40 | NA | E3 | Remission | 1 | NA | Past | No | No | No |
| CD3 | 2020 | Female | 38 | NA | L3L4 | Flare | 1 | 4 | Never | No | No | No |
| CD4 | 2020 | Male | 44 | NA | L3L4 | Remission | 0.6 | 1 | Never | No | No | No |
| CD5 | 13/08/2020 | Female | 40 | 20 | L3 | Remission | 5 | 2 | Past | No | No | Yes |
| CD6 | 15/09/2020 | Male | 60 | 55 | L3 | Flare | 5 | 7 | Never | No | No | Yes |
| CD7 | 24/05/2019 | Female | 26 | 16 | L3 | Flare | 89.9 | NA | Past | No | No | No |
| CD8 | 10/07/2020 | Male | 42 | 13 | L3 | Flare | 6.3 | 3 | Never | No | No | Yes |
| CD9 | 06/07/2020 | Male | 47 | 12 | L2 | Remission | 7.8 | 0 | Past | No | No | No |
| CD10 | 31/05/2018 | Female | 50 | 47 | L1 | Remission | 1 | NA | Current | No | No | Yes |
| CD11 | 21/06/2020 | Female | 33 | 19 | L3 | Flare | 5.3 | NA | Past | No | No | No |
| CD12 | 03/07/2020 | Male | 22 | 20 | L3 | Remission | 5 | 3 | Never | No | No | Yes |
| CD13 | 16/06/2020 | Male | 56 | 52 | L1 | Flare | 8.7 | 7 | Past | No | No | Yes |
| CD14 | 26/10/2020 | Female | 56 | 20 | L3 | Remission | 1 | 2 | Never | No | No | Yes |
| CD15 | 02/10/2020 | Female | 37 | 27 | L3 | Flare | 1 | 7 | Past | No | No | No |
| H1 | 2020 | Female | 26 | NA | NA | NA | NA | NA | Never | No | No | No |
| H2 | 2020 | Female | 28 | NA | NA | NA | NA | NA | Never | No | No | No |
| H3 | 2020 | Male | 31 | NA | NA | NA | NA | NA | Never | No | No | No |
| H4 | 2020 | Male | 29 | NA | NA | NA | NA | NA | Never | No | No | No |
| H5 | 2020 | Male | 36 | NA | NA | NA | NA | NA | Never | No | No | No |
| H6 | 2021 | Female | 25 | NA | NA | NA | NA | NA | Current | No | No | No |
| H7 | 2021 | Male | 32 | NA | NA | NA | NA | NA | Never | No | No | No |
| H8 | 2021 | Female | 38 | NA | NA | NA | NA | NA | Current | No | No | No |
| H9 | 2021 | Female | 27 | NA | NA | NA | NA | NA | Never | No | No | No |
| H10 | 2021 | Female | 29 | NA | NA | NA | NA | NA | Never | No | No | No |
| H11 | 2021 | Female | 56 | NA | NA | NA | NA | NA | Never | No | No | No |
| H12 | 2021 | Female | 41 | NA | NA | NA | NA | NA | Never | No | No | No |
| H13 | 2021 | Female | 28 | NA | NA | NA | NA | NA | Never | No | No | No |
| H14 | 2021 | Female | 55 | NA | NA | NA | NA | NA | Never | No | No | No |

**Table S1: Detailed information about the individuals included in the study.**

| SampleName | Status | Sample Type | Raw | Clean and non-human |
|---|---|---|---|---|
| H1_F | Healthy | Feces | 4 557 777 | 3 302 598 |
| H2_F | Healthy | Feces | 5 141 556 | 3 538 237 |
| H3_F | Healthy | Feces | 4 575 463 | 3 314 041 |
| H4_F | Healthy | Feces | 5 006 463 | 3 702 841 |
| H5_F | Healthy | Feces | 6 630 408 | 4 868 696 |
| H6_F | Healthy | Feces | 6 043 597 | 4 533 278 |
| H7_F | Healthy | Feces | 149 | 113 |
| H8_F | Healthy | Feces | 12 607 586 | 10 702 899 |
| H9_F | Healthy | Feces | 4 957 375 | 4 008 417 |
| H10_F | Healthy | Feces | 161 | 122 |
| H11_F | Healthy | Feces | 47 423 047 | 36 124 946 |
| H12_F | Healthy | Feces | 3 848 503 | 3 021 191 |
| H13_F | Healthy | Feces | 3 944 094 | 3 101 396 |
| H14_F | Healthy | Feces | 10 220 371 | 8 099 646 |
| CD1_F | Crohn | Feces | 3 398 455 | 2 610 231 |
| CD2_F | Crohn | Feces | 11 438 302 | 8 205 616 |
| CD3_F | Crohn | Feces | 27 092 512 | 19 524 928 |
| CD4_F | Crohn | Feces | 18 270 751 | 12 128 024 |
| CD5_F | Crohn | Feces | 8 453 588 | 5 729 131 |
| CD6_F | Crohn | Feces | 4 522 582 | 3 789 808 |
| CD7_F | Crohn | Feces | 2 985 683 | 2 478 116 |
| CD8_F | Crohn | Feces | 10 219 909 | 8 279 831 |
| CD9_F | Crohn | Feces | 7 091 001 | 5 360 552 |
| CD10_F | Crohn | Feces | 9 143 203 | 7 378 428 |
| CD11_F | Crohn | Feces | 9 290 609 | 7 084 801 |
| CD12_F | Crohn | Feces | 6 350 692 | 5 222 735 |
| CD13_F | Crohn | Feces | 7 983 876 | 6 487 386 |
| CD14_F | Crohn | Feces | 636 447 | 498 993 |
| CD15_F | Crohn | Feces | 3 500 715 | 2 724 997 |
| H1_B | Healthy | Blood | 25 145 743 | 6 790 645 |
| H2_B | Healthy | Blood | 13 872 445 | 5 336 467 |
| H3_B | Healthy | Blood | 17 469 863 | 6 122 420 |
| H4_B | Healthy | Blood | 16 038 090 | 3 433 116 |
| H5_B | Healthy | Blood | 19 342 807 | 5 840 315 |
| H6_B | Healthy | Blood | 1 916 528 | 185 563 |
| H7_B | Healthy | Blood | 2 347 924 | 837 299 |
| H8_B | Healthy | Blood | 2 733 222 | 788 996 |
| H9_B | Healthy | Blood | 2 689 347 | 723 438 |
| H10_B | Healthy | Blood | 2 486 126 | 573 092 |
| H11_B | Healthy | Blood | 1 397 921 | 160 240 |
| H12_B | Healthy | Blood | 3 711 044 | 691 480 |
| H13_B | Healthy | Blood | 2 312 559 | 68 548 |
| H14_B | Healthy | Blood | 5 506 340 | 2 713 132 |
| CD1_B | Crohn | Blood | 25 941 062 | 8 696 199 |
| CD2_B | Crohn | Blood | 14 183 825 | 8 720 837 |
| CD3_B | Crohn | Blood | 29 669 303 | 14 833 794 |
| CD4_B | Crohn | Blood | 20 129 214 | 11 955 938 |
| CD5_B | Crohn | Blood | 16 967 073 | 10 236 927 |
| CD6_B | Crohn | Blood | 4 631 975 | 1 917 021 |
| CD7_B | Crohn | Blood | 2 048 220 | 12 333 |
| CD8_B | Crohn | Blood | 7 197 457 | 3 434 820 |
| CD9_B | Crohn | Blood | 6 813 381 | 3 293 988 |
| CD10_B | Crohn | Blood | 6 542 891 | 3 338 429 |
| CD11_B | Crohn | Blood | 7 374 010 | 2 778 364 |
| CD12_B | Crohn | Blood | 9 614 014 | 4 117 958 |
| CD13_B | Crohn | Blood | 5 606 138 | 2 207 758 |
| CD14_B | Crohn | Blood | 5 555 652 | 2 052 908 |
| CD15_B | Crohn | Blood | 8 436 207 | 4 380 873 |
| Microtube | | | 5 242 956 | 2 930 415 |
| BloodCollectionTube | | | 10 853 654 | 4 427 872 |

**Table S2**: **Number of read pairs per sample.**
The number of pairs obtained after sequencing is indicated in the "Raw" column. The number of pairs remaining after quality-filtering and removal of non-human reads is in indicated in the "Clean and non-human" column. The samples with less than 100 000 pairs of clean and non-human reads, highlighted in red, were excluded from the analysis.

**Table S3: Characteristics of all the vOTUs.**

The table includes quality, length, annotation, classification, host prediction, homology with oral and gut viral databases for each vOTU. The table is organized as follows: first the non-contaminant vOTUs, those found in blood samples then in fecal samples, and then the contaminant vOTUs (no particular order).

*Table S3 is too large and is available as a separate Excel file.*
